# Network medicine links SARS-CoV-2/COVID-19 infection to brain microvascular injury and neuroinflammation in dementia-like cognitive impairment

**DOI:** 10.1101/2021.03.15.435423

**Authors:** Yadi Zhou, Jielin Xu, Yuan Hou, James B. Leverenz, Asha Kallianpur, Reena Mehra, Yunlong Liu, Haiyuan Yu, Andrew A. Pieper, Lara Jehi, Feixiong Cheng

## Abstract

**Background:** Dementia-like cognitive impairment is an increasingly reported complication of SARS-CoV-2 infection. However, the underlying mechanisms responsible for this complication remain unclear. A better understanding of causative processes by which COVID-19 may lead to cognitive impairment is essential for developing preventive interventions.

**Methods:** In this study, we conducted a network-based, multimodal genomics comparison of COVID-19 and neurologic complications. We constructed the SARS-CoV-2 virus-host interactome from protein-protein interaction assay and CRISPR-Cas9 based genetic assay results, and compared network-based relationships therein with those of known neurological manifestations using network proximity measures. We also investigated the transcriptomic profiles (including single-cell/nuclei RNA-sequencing) of Alzheimer’s disease (AD) marker genes from patients infected with COVID-19, as well as the prevalence of SARS-CoV-2 entry factors in the brains of AD patients not infected with SARS-CoV-2.

**Results:** We found significant network-based relationships between COVID-19 and neuroinflammation and brain microvascular injury pathways and processes which are implicated in AD. We also detected aberrant expression of AD biomarkers in the cerebrospinal fluid and blood of patients with COVID-19. While transcriptomic analyses showed relatively low expression of SARS-CoV-2 entry factors in human brain, neuroinflammatory changes were pronounced. In addition, single-nucleus transcriptomic analyses showed that expression of SARS-CoV-2 host factors (*BSG* and *FURIN*) and antiviral defense genes (*LY6E*, *IFITM2*, *IFITM3*, and *IFNAR1*) was significantly elevated in brain endothelial cells of AD patients and healthy controls relative to neurons and other cell types, suggesting a possible role for brain microvascular injury in COVID-19-mediated cognitive impairment. Notably, individuals with the AD risk allele *APOE* E4/E4 displayed reduced levels of antiviral defense genes compared to *APOE* E3/E3 individuals.

**Conclusion:** Our results suggest significant mechanistic overlap between AD and COVID-19, strongly centered on neuroinflammation and microvascular injury. These results help improve our understanding of COVID-19-associated neurological manifestations and provide guidance for future development of preventive or treatment interventions.

## Introduction

Patients with COVID-19 commonly develop neurologic symptoms and/or complications, such as a loss of taste or smell, stroke, delirium, and rarely new onset seizures [1, 2]. Based on the experience with other coronaviruses, it was predicted early on that COVID-19 patients might also be at risk for cognitive dysfunction. For example, after the severe acute respiratory syndrome (SARS-CoV-1) outbreak in 2002 and the Middle East respiratory syndrome (MERS) outbreak in 2012, both caused by human coronaviruses (HCoVs), 20% of recovered patients reported ongoing memory impairment [3]. Evidence now supports similar complications after COVID-19, which due to the global pandemic, is poised to potentially lead to a surge in cases of Alzheimer’s-like dementia or other forms of neurocognitive impairment in the near future [4, 5].

Clarification of the underlying molecular mechanisms of COVID-19-induced cognitive impairment is mandatory for developing effective therapeutic strategies for patients [6–8]. While some studies have shown that SARS-CoV-2 may directly infect the brain [9–11], potentially through the olfactory bulb [9], others have shown that SARS-CoV-2 is absent from the brain [12] and cerebrospinal fluid (CSF) [13]. COVID-19 has also been suggested to cause inflammation within the central nervous system (CNS) [8, 12, 14], as well as microvascular injury [12]. For example, the SARS-CoV-2 spike protein, which readily crosses the blood-brain barrier (BBB) [15, 16], induces an inflammatory response within microvascular endothelial cells, leading to BBB dysfunction [16].

Multi-omics datasets for patients with COVID-19, such as bulk and single-cell/nucleus transcriptomic [17], proteomic [18], and interactomic (protein-protein interactions [PPIs]) datasets [19–23], have been generated in order to conduct unbiased investigation of the pathophysiological pathways. We reasoned that network-based drug-disease and disease-disease proximity approaches [24–27], which shed light on the relationship between drugs (and drug targets) and diseases (gene and protein determinants of disease mechanisms in the human PPI network), would provide mechanistic insights into the pathobiology of cognitive dysfunction after SARS-CoV-2 infection, potentially suggesting novel targets for further therapeutic investigation. Thus, we investigated Alzheimer’s disease (AD)-like pathobiology associated with SARS-CoV-2 infection by using a network-based multimodal omics analytic methodology (**Fig. 1**). Specifically, we leveraged bulk and single-cell/nuclei RNA-sequencing, proteomics, and interactomics (SARS-CoV-2 virus-host PPIs from mass spectrometry assays and genetic interactions from CRISPR-Cas9 assays) from COVID-19 and AD patients. We hypothesized that SARS-CoV-2 host factors would be localized in a subnetwork within the comprehensive PPI network and that proteins associated with certain neurologic function would be targeted by the virus either directly, or indirectly through PPIs with virus host factors. As detailed below, our comprehensive analyses show scant evidence of direct brain and neuron damage by COVID-19, but robust evidence for involvement of pathways of neuroinflammation and brain microvascular injury in COVID-19.

**Fig. 1.**
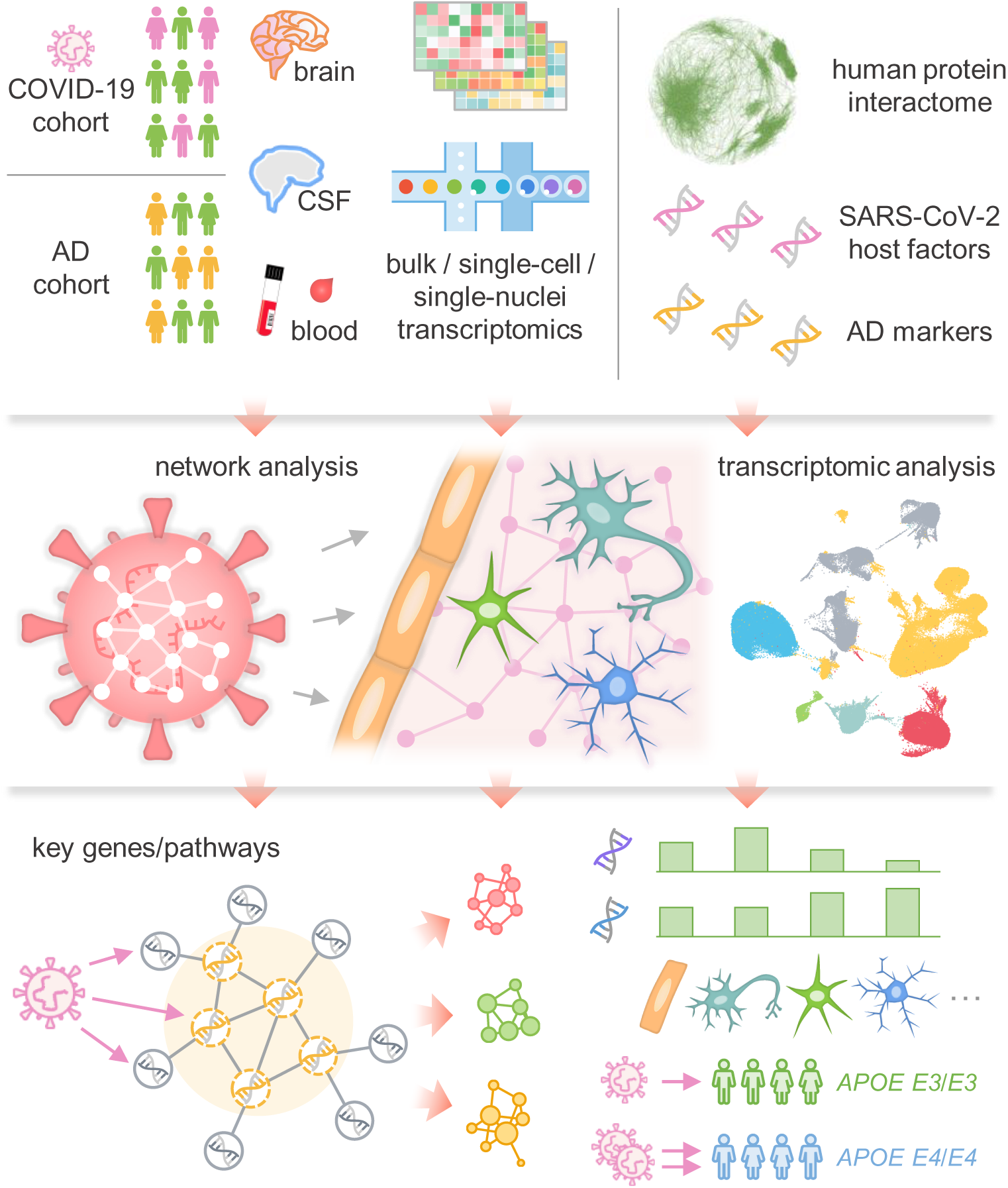
Overall workflow of this study. We compiled ten SARS-CoV-2 host factor datasets based on CRISPR-Cas9 assays or protein-protein interaction assays, and collected neurological disease-associated genes/proteins. We utilize network proximity analysis to investigate network-based relationship between SARS-CoV-2 host factors and neurological disease-associated genes/proteins under the human interactome network model. Utilizing bulk/single-cell/single-nuclei transcriptomics data, AD markers, and SARS-CoV-2 entry factors, we tested three potential mechanisms of SARS-CoV-2 neurological manifestations: direct brain invasion, neuroinflammation, and microvascular injury. The susceptibility of SARS-CoV-2 infection was also compared among AD patients with different *APOE* genotypes.

## Materials and methods

### SARS-CoV-2 host factor profiles

In total, we have gathered ten datasets of SARS-CoV-2 (and other HCoVs) target host genes/proteins from various data sources (**Table S1**). Specifically, six of these datasets were based on CRISPR-Cas9 assay results, including (1-2) CRISPR_A549-H and CRISPR_A549-L, based on high (-H) and low (-L) multiplicity of infection of SARS-CoV-2 in A549 cells [21]; (3-5) CRISPR_HuH7-SARS2, CRISPR_HuH7-229E, CRISPR_HuH7-OC43, based on HuH7 cells infected by SARS-CoV-2, HCoV-229E, and HCoV-OC43, respectively [22]; and (6) CRISPR_VeroE6, based on SARS-CoV-2-infected VeroE6 cells [23]. For the CRISPR-Cas9-based datasets, we considered the top-100 host factors using the ranking methods described in the respective original publications [21–23]. We also examined the effect of using top-50, −150, and −200 genes. In addition to the CRISPR datasets, we collected three mass spectrometry-based virus-host PPI datasets [19, 20] for SARS-CoV-2, SARS-CoV-1, and MERS-CoV, named as SARS2-PPI, SARS1-PPI, and MERS-PPI. The last dataset, HCoV-PPI, was from our recent studies [28, 29] containing HCoVs target host proteins supported by literature-based evidence. Functional enrichment analyses, including Kyoto Encyclopedia of Genes and Genomes (KEGG) and Gene Ontology (GO) biological process enrichment analyses, were performed using Enrichr [30] for the CRISPR datasets. A list of main SARS-CoV-2 entry factors and proteins involved in antiviral defense was assembled [8], including *ACE2*, *BSG*, *NRP1*, *TMPRSS2*, *TMPRSS11A*, *TMPRSS11B*, *FURIN*, *CTSB*, *CTSL*, *LY6E*, *IFITM1*, *IFITM2*, *IFITM3*, *IFNAR1*, and *IFNAR2*.

### Neurological disease gene profiles

We extracted neurologic disease-associated genes/proteins from the Human Gene Mutation Database (HGMD) [31], and defined a gene to be disease-associated, if it had at least one disease-associated mutation from HGMD reported in the literature. The details of these neurological disease genes can be found in **Table S2**, including the reported mutations, disease terms used to identify the neurological diseases [32], and original references. For AD, we assembled four datasets from AlzGPS (https://alzgps.lerner.ccf.org/) [33], based on our previous work [34] (**Table S2**). These datasets contain experimentally validated genes (denoted as “seed” genes) in amyloid pathology (amyloid) or tauopathy (tau), as well as high-confidence AD risk genes identified by genome-wide association study (GWAS) [35].

### Alzheimer’s disease blood and CSF markers

We compiled a list of AD blood and CSF protein markers from previous studies [36–38], which included 29 blood markers and 31 CSF markers. The expression alteration of these markers in AD or AD-related pathologies, such as tauopathy, were extracted from these studies. The details of these markers can be found in **Table S3**.

### Transcriptomic data analyses

Two categories of transcriptomic datasets, including three from AD patients and three from COVID-19 patients, were used (**Table S4**). These datasets are described below. All single-cell analyses were performed using Seurat v3.1.5 [39] following the processing steps from the original publication of each dataset. Cell types were identified using markers based on the original publications, unless already annotated in the metadata. Differential expression analysis was performed using the “FindMarkers” function from Seurat for the single-cell/nuclei datasets. For the bulk RNA-sequencing dataset, differential expression analysis was performed using edgeR v3.12 [40]. Differentially expressed genes (DEGs) were determined by false discovery rate (FDR) < 0.05 and |log_2_foldchange| > 0.5.

#### GSE147528

This single-nuclei RNA-sequencing dataset from the superior frontal gyrus and entorhinal cortex regions of 10 males with varying stages of AD [41] was used to examine the expression of the four key SARS-CoV-2 entry factors: *ACE2*, *TMPRSS2*, *FURIN*, and *NRP1,* in neurons.

#### GSE157827

This single-nuclei RNA-sequencing dataset from the prefrontal cortex region of 12 AD patients and 9 normal controls [42] was used to test the susceptibility of brain endothelial cells to SARS-CoV-2 infection and damage. Six cell types were included: astrocytes, endothelial cells, excitatory neurons, inhibitory neurons, microglia, and oligodendrocytes. The *APOE* genotypes of these individuals are also available in this dataset.

#### GSE138852

This single-nuclei RNA-sequencing dataset from the entorhinal cortex of individuals with AD (n = 6) and healthy controls (n = 6) [43] was used to validate the findings of the expression of SARS-CoV-2 entry factors in brain endothelial cells. Six cell types were included: astrocytes, endothelial cells, neurons, microglia, oligodendrocytes, and oligodendrocyte progenitor cells.

#### GSE157103

This bulk RNA-sequencing dataset of 125 peripheral blood mononuclear cell (PBMC) samples [44] was used to examine the expression spectrum of AD blood biomarkers. DEGs were analyzed by disease severity conditions: 66 intensive care unit (ICU) patients (COVID-19 patients n = 50 vs. non-COVID-19 patients n = 16), 59 non-ICU patients (COVID-19 patients n = 49 vs. non-COVID-19 patients n = 10), and all 125 patients. Adjustments for the effects of age and sex were made.

#### GSE149689

This single-cell RNA-sequencing PBMC dataset of 6 samples from severe COVID-19 patients, 4 samples from mild COVID-19 patients, and 4 samples from healthy controls [45] was used to examine the expression spectrum of AD blood markers. 13 cell types were included in this dataset: lgG^-^ B cells, lgG^+^ B cells, CD4^+^ T cell effector memory (EM)-like cells, CD4^+^ T cell non-EM-like cells, CD8^+^ T cell EM-like cells, CD8^+^ T cell non-EM-like cells, dendritic cells, monocytes, intermediate monocytes, nonclassical monocytes, natural killer cells, platelets, and red blood cells.

#### GSE163005

This single-cell RNA-sequencing CSF dataset [46] was used to examine the expression spectrum of AD CSF markers. This neuro-COVID-19 dataset contains 8 COVID-19 patients, 9 multiple sclerosis (MS) patients, 9 idiopathic intracranial hypertension (IIH) patients, and 5 viral encephalitis (VE) patients. Based on the original publication, the cells were categorized into three major cell groups of T cells, dendritic cells, and monocytes. Four comparisons were performed for each major cell group: COVID-19 vs. MS, COVID-19 vs. IIH, COVID-19 vs. VE, and COVID-19 vs. non-COVID-19 (MS, IIH, and VE).

### Human protein-protein interactome

The human protein-protein interactome was from our previous studies [24, 25, 47, 48], and contains 17,706 protein nodes and 351,444 unique PPI edges. Each PPI edge has one or more source information of five categories of evidence from publicly available databases and datasets: protein complexes identified by robust affinity purification-mass spectrometry from BioPlex V2.016 [49]; binary PPIs discovered by high-throughput yeast two-hybrid systems in three datasets [24, 50, 51]; signaling networks revealed by low-throughput experiments from SignaLink2.0 [52]; low-throughput or high-throughput experiments uncovered kinase-substrate interactions from KinomeNetworkX [53], Human Protein Resource Database (HPRD) [54], PhosphoNetworks [55], PhosphositePlus [56], DbPTM 3.0 [57], and Phospho.ELM [58]; and PPIs curated from literatures identified by yeast two-hybrid studies, affinity purification-mass spectrometry, low-throughput experiments, or protein three-dimensional structures from BioGRID [59], PINA [60], Instruct [61], MINT [62], IntAct [63], and InnateDB [64]. Inferred PPIs derived from evolutionary analysis, gene expression data, and metabolic associations were excluded.

### Network analyses

We used network proximity metrics to quantify the network associations of two gene/protein modules. The “shortest” proximity measure was used to evaluate the overall average distance among all genes in the neurological disease gene sets and the SARS-CoV-2 host factor profiles:

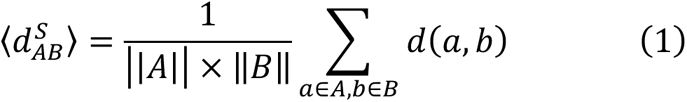

where *d*(*a*, *b*) represents the shortest path length between gene *a* from module *A* and *b* from module *B* in the human protein-protein interactome. “closest” proximity measure was used to quantify the distance among the AD markers and the DEGs from the COVID-19 omics datasets focusing on the genes that are closest to the genes in the other module:

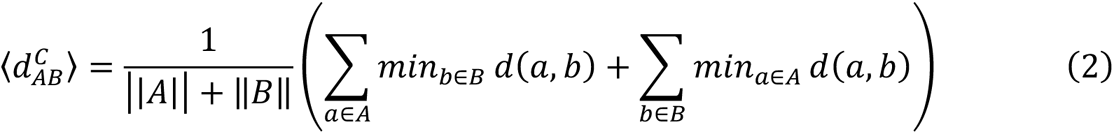

All network proximities were converted to Z scores based on permutation tests of 1000 repeats:

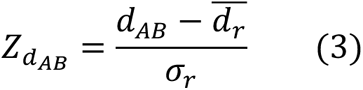

where 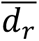 and *σ*_F_ are the mean and standard deviation of the proximities, respectively. A P value was computed using the permutation test accordingly. Gene set pairs with P < 0.05 and Z < −1.5 were considered significantly proximal.

The largest connect component (LCC) was computed by NetworkX [65]. Significance of LCC was computed in the same way as the network proximity using permutation test repeated 1000 times. Eigenvector centrality [66] of the nodes in the networks were computed using Gephi 0.9.2 [67] to evaluate the influence of the nodes considering the importance of their neighbors.

### Tissue and brain region expression specificity

We retrieved the transcriptomic data in raw count and transcripts per million (TPM) from the GTEx v8 release [68] for 33 human tissues and 13 brain regions, and examined expression across different tissues and brain regions. At the tissue level, the brain regions were combined as one “brain” tissue. We first defined a gene to be tissue-or brain region-expressed if it had a count per million (CPM) ≥ 0.5 in over 90% samples. Then, to quantify the significance of the expression of a gene in a tissue or brain region, we normalized its expression using the z score method.

### Innate immune genes

We retrieved a list of 1031 human innate immunity genes from InnateDB [64], which were associated in the published literature with roles in innate immunity.

### Statistical analysis and network visualization

Python package SciPy v1.3.0 [69] was used for the statistical tests unless specified otherwise. *P* < 0.05 (or FDR < 0.05 when applicable) was considered statistically significant throughout the study. Networks were visualized with Gephi 0.9.2 [67] and Cytoscape 3.8.0 [70].

## Results

### A network-based, multimodal omics analytic framework

In this study, we present a network-based, multimodal omics (including bulk and single-cell/nuclei RNA-sequencing, proteomics, and interactomics) analysis method for investigating the underlying mechanisms of COVID-19-associated cognitive dysfunction or impairment. We hypothesized that for COVID-19 to have neurological impacts in the host CNS, its host factors (genes/proteins) should be localized in the corresponding subnetwork within the human PPI network, and either directly target the neurological disease-associated genes/proteins or indirectly affect them through PPIs (**Fig. 1**). We utilized single-cell/nuclei RNA-sequencing data from both COVID-19 patients with neurological manifestations (neuro-COVID-19) and brains of AD patients not infected by SARS-CoV-2, brain-region specific gene expression data from the GTEx database [68], SARS-CoV-2 virus-host PPIs from mass spectrometry assays, genetic interactions from CRISPR-Cas9 assays (**Table S1**), and disease-related genetic data (**Table S2**).

We compiled ten virus-host interaction datasets across SARS-CoV-2, SARS-CoV-1 and MERS-CoV, and other common HCoVs, including six datasets from CRISPR-Cas9 assays and four datasets for virus-human PPIs (**Table S1**). Functional enrichment analyses of each dataset revealed that virus-host PPIs and host factors are significantly enriched in pathways well-known to be involved in SARS-CoV-2 infection and related immune responses (**Supplementary Results**, **Fig. S1**). Using these datasets, we computed their network associations with ten neurological diseases or conditions. To determine whether brain damage was caused by SARS-CoV-2 direct infection of the brain, we evaluated expression levels of SARS-CoV-2 entry genes at brain region and brain single-cell levels. Neuroinflammation was evaluated by identifying alterations in expression of AD blood and CSF biomarkers in COVID-19 patients using data from peripheral blood mononuclear cell (PBMC) and CSF samples (neuro-COVID-19 dataset). Lastly, microvascular injury was evaluated by examining the expression of SARS-CoV-2 entry factors and antiviral defense genes in brain endothelial cells of AD and healthy control samples. We also compared the expression of SARS-CoV-2 entry factors and antiviral defense genes in individuals with different *APOE* genotypes.

### Strong network-based relationships of COVID-19 to neurological manifestations

We assembled experimentally validated gene/protein profiles for ten neurological diseases or conditions, including AD, amyotrophic lateral sclerosis, cognitive decline, dementia, frontotemporal dementia, multiple system atrophy, neuronal ceroid lipofuscinosis, Parkinson’s disease (PD), spinal muscular atrophy, and spinocerebellar ataxia (**Table S2**). First, we quantified the network distance of the SARS-CoV-2 host factor datasets and neurological diseases in the human protein-protein interactome. A close network distance between SARS-CoV-2 host factors and neurological disease-associated genes/proteins suggests related or shared mechanistic pathways between COVID-19 and specific neurological disease [29]. Using state-of-the-art network proximity measures (see Methods), we evaluated the network-based relationship for the gene/protein sets between virus-host factors and each disease/condition under the human interactome network model (**Fig. 2a** and **Fig. S2**). We found significant proximities between the SARS-CoV-2 virus-host interactome (including PPIs and genetic interactions) and genes associated with neurological diseases in the human interactome network (average Z = −1.82). The SARS-CoV-2 virus-host PPIs (average Z = −2.54) showed more significant network proximities (white circles, **Fig. 2a**) compared to CRISPR-Cas9-derived host factors (average Z = −1.34). The top three neurological diseases or conditions with the smallest network proximities to SARS-CoV-2 were: AD (average Z = −2.75) [6, 7], cognitive decline (average Z = −2.77), and PD (average Z = - 2.94). Recent case reports of COVID-19 patients developing parkinsonism suggest that COVID-19 patients may have increased risk of PD later in life [71]. We noticed that amyloid pathology has significant network proximity (average Z = −1.55) with the PPI datasets. However, there are no significant network-based relations between tauopathy-related genes and the SARS-CoV-2 interactome. One possible explanation is the incompleteness of genes/proteins related to tauopathy in the datasets. In addition to SARS-CoV-2, HCoV-229E also showed a significant network proximity to neurological diseases, suggesting a common association between coronaviruses and cognitive dysfunction [72].

**Fig. 2.**
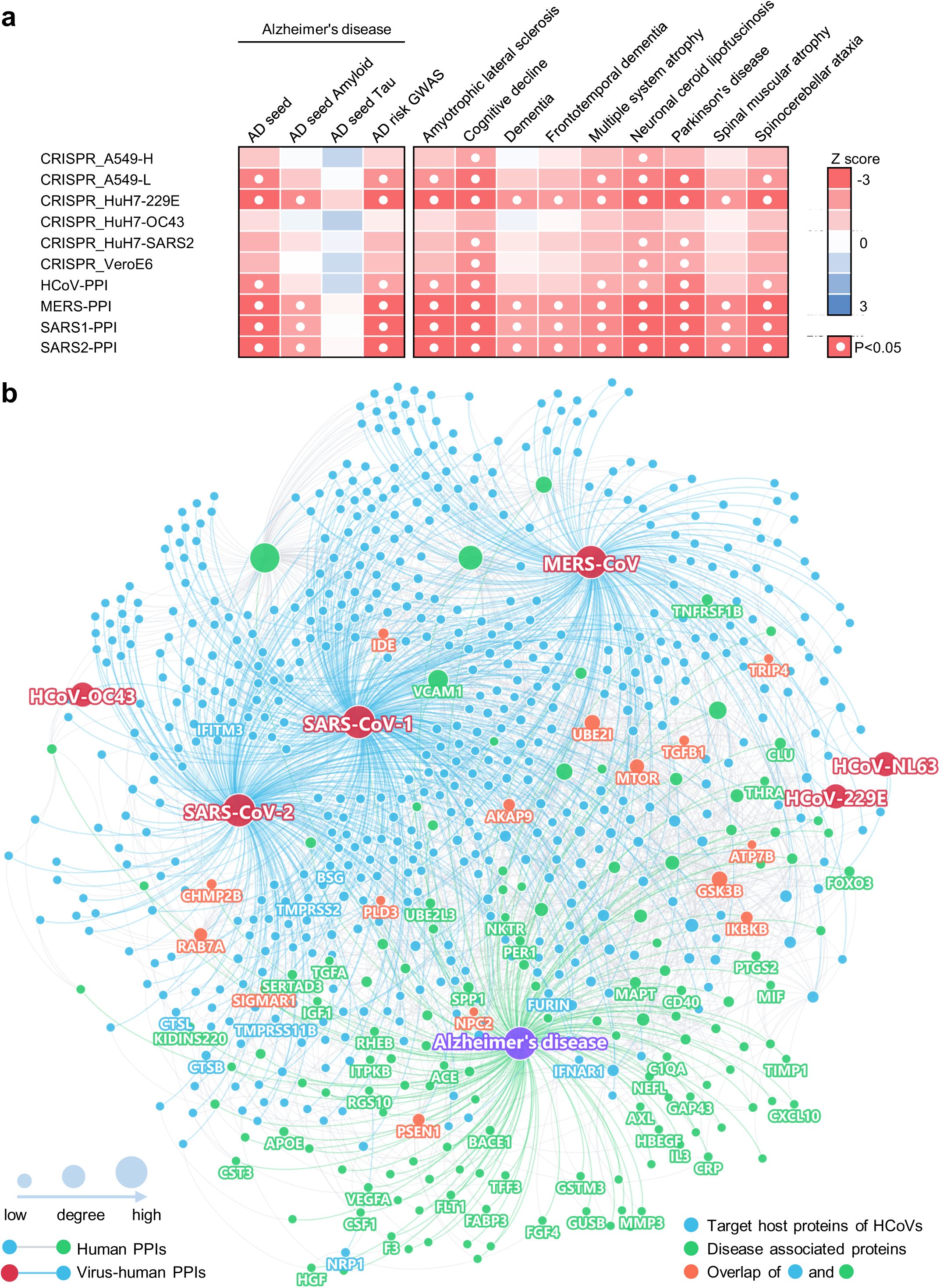
A network landscape of COVID-19 and neurological diseases. (**a**) Network proximity analysis shows strong network associations between COVID-19 and neurological diseases. Heatmap shows the “shortest” network proximities in Z score (see Methods). Smaller Z scores indicate smaller network proximities between the two gene sets. (**b**) Protein-protein interaction network of the SARS-CoV-2 and other human coronaviruses host factors and the Alzheimer’s disease-associated genes/proteins. SARS-CoV-2 entry factors, antiviral defense genes, and AD biomarkers are highlighted by their gene symbols.

### A network-based relationship between COVID-19 and Alzheimer’s disease

To examine further why cognitive impairment has such significant network-based association with the SARS-CoV-2 interactome, we focused on AD and visualized the PPIs among AD seed genes/proteins (**Fig. 2b**, green nodes) and host genes/proteins illustrated by the four SARS-CoV-2 virus-human PPI datasets (**Fig. 2b**, blue nodes). We found a large number of PPIs among these proteins, including multiple blood and CSF biomarkers and SARS-CoV-2 entry factors (nodes with gene symbols). Here, we discuss several markers that may have important roles in COVID-19-associated AD (**Table S5**) according to network measures (connectivity and eigenvector centrality [EC]), including vascular cell adhesion protein 1 (VCAM1) (connectivity K = 73), ras-related protein Rab-7a (RAB7A) (K = 30), and transforming growth factor beta 1 (TGFB1) (K = 10). These proteins also have high EC values, a measure of potential node (gene/protein) influence on the network that considers the influence of its neighbors: VCAM1 EC = 0.59 (rank 6 out of 153 AD genes/proteins), RAB7A EC = 0.17 (rank 25), and TGFB1 EC = 0.19 (rank 22).

VCAM1 is located at the endothelial cell surface and is activated by cytokines [73]. It is also an AD biomarker with elevated expression in the blood [74, 75] and CSF [36, 37] of AD patients. VCAM1 levels were also significantly associated with the severity of dementia and structure changes of white matter [75], and brain endothelial VCAM1 at the blood-brain barrier has been proposed as a target for treating age-related neurodegeneration [76]. Serum VCAM1 levels were also significantly elevated in severe COVID-19 patients compared to mild patients and controls, and significantly decreased in the convalescence phase compared to severe patients [77]. Notably, VCAM1 also plays an important role in COVID-19-induced vasculitis [78]. RAB7A is a direct target of non-structural protein 7 (nsp7) of SARS-CoV-2 [20], and also one of the top host factors in CRISPR-Cas9-based SARS-CoV-2 datasets. RAB7A knockout reduces cell surface angiotensin converting enzyme 2 (ACE2) levels, which thereby reduces SARS-CoV-2 entry into cells [21]. RAB7A is also a potential AD biomarker whose blood expression level is positively associated with high memory test performance [38]. TGFB1 is a cytokine that controls cell growth and differentiation [79, 80] and a potential AD marker with decreased expression in the blood of AD patients [38]. The anti-inflammatory and neuroprotective role of TGFB1 against AD has already been demonstrated in animal models [81, 82]. Using bulk RNA-sequencing data from PBMC samples of COVID-19 patients, we also found that TGFB1 expression was significantly decreased in both mild COVID-19 patients and those requiring intensive care unit (ICU) level care, as compared to non-COVID-19 patients (**Table S3**).

Altogether, these results encouraged us to explore further the pathological relationships between COVID-19 and AD, and to identify potential pathological pathways by which SARS-CoV-2 infection could lead to AD-like dementia.

### Neuroinflammation-mediated association between neuro-COVID-19 and AD

We next turned to investigate whether neuroinflammation was a shared mechanism between COVID-19 and AD by investigating the expression levels of well-known AD blood and CSF marker genes in COVID-19 patients with neurological manifestations (neuro-COVID-19). To this end, we compiled a list of blood and CSF protein markers for AD from previous studies [36–38] (**Table S3**) with their expression alterations in AD or AD-related pathologies. We then examined their expression in COVID-19 patient PBMC [44, 45] and CSF [46] samples. We performed differential expression analyses for the PBMC bulk RNA-sequencing dataset [44] of COVID-19 patients vs. non-COVID-19 patients. For the other single-cell level PBMC dataset [45], we compared mild / severe COVID-19 patients to healthy controls. We used an additional single-cell RNA-sequencing dataset generated from CSF samples of neuro-COVID-19 patients with well-defined neurological manifestations [46].

We first examined the degree of overlap between AD markers and differentially expressed genes (DEGs) in PBMCs or CSF from COVID-19 patients and found significant overlap in CSF monocytes (p = 0.004, Fisher’s exact test, **Table S3**), but not in PBMCs (p = 0.807, **Table S3**). We further computed the network proximities of the AD markers and DEGs and found that blood markers and DEGs from PBMCs do not show significant network proximities, whereas CSF markers and DEGs from CSF monocytes were significantly proximal (**Table S3**, Z = −3.69, p = 0.002). Altogether, we found a more significant network-based relationship between COVID-19 and AD in CSF (including monocytes) compared to PBMCs from COVID-19 patients. We next examined the overall expression spectrum of these markers in both PBMCs and CSF (**Fig. 3a-b**).

**Fig. 3.**
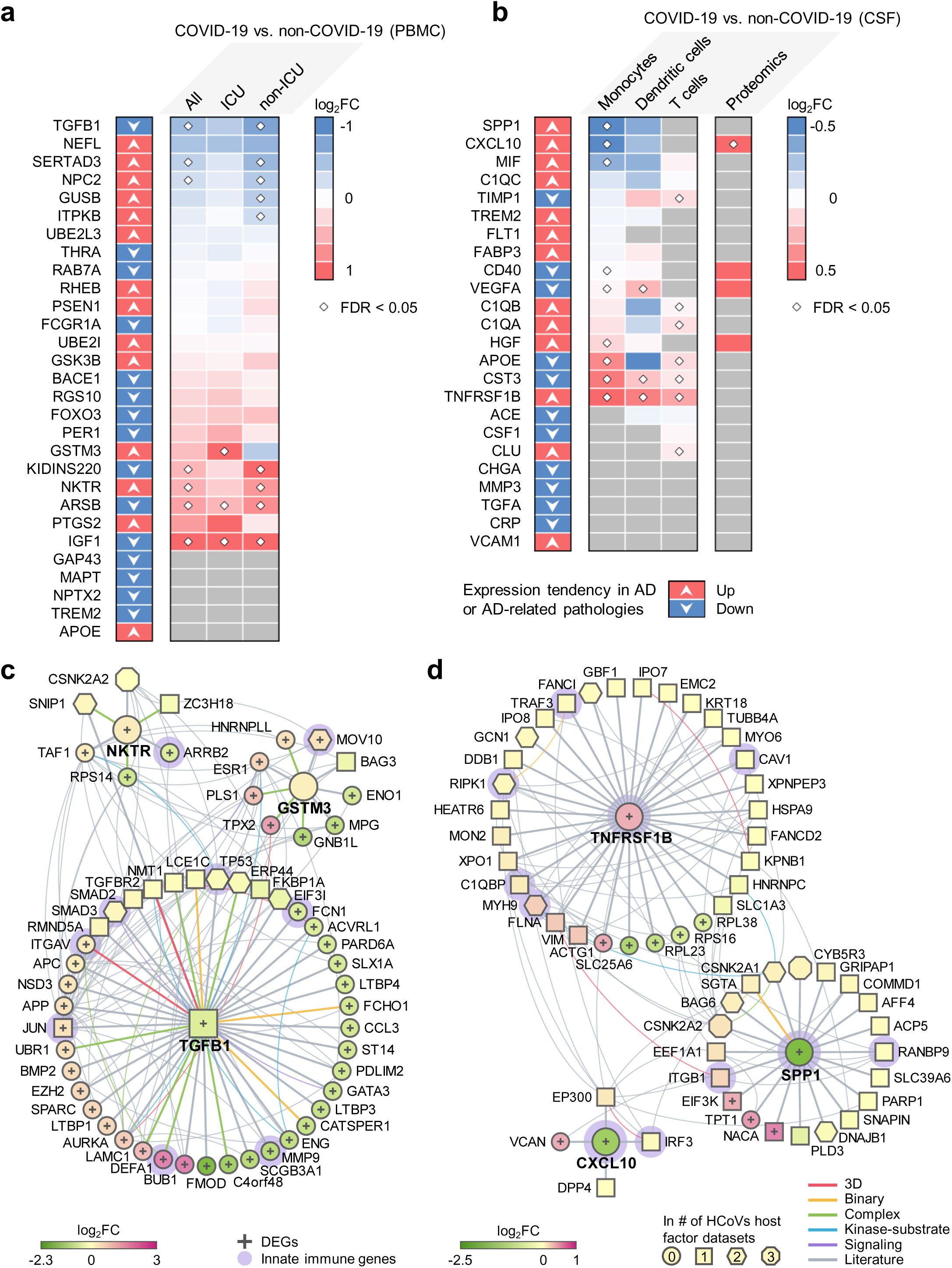
Neuroinflammation-mediated association between COVID-19 and Alzheimer’s disease (AD). The expression of (**a**) AD blood and (**b**) cerebrospinal fluids (CSF) protein markers in COVID-19 patients. Heatmaps show the fold change (FC) of the comparisons indicated above. (**c**) and (**d**) Network analyses of the AD markers that are differentially expressed in COVID-19 vs. non-COVID-19. Neighbors of these markers that are the SARS-CoV-2 host factors (non-circle nodes) or are DEGs (denoted by “+”) in the COVID-19 datasets are shown. Node shape indicates the number of SARS-CoV-2 host factor datasets that contain the node. Edge colors indicate the protein-protein interaction source type. PBMC, peripheral blood mononuclear cells. DEG, differentially expressed genes.

In PBMCs, the expression of several AD markers was altered by SARS-CoV-2 infection, such as *TGFB1*, SERTA domain-containing protein 3 (*SERTAD3*), glutathione S-transferase M3 (*GSTM3*), kinase D-interacting substrate of 220 kDa (*KIDINS220*), natural killer tumor recognition sequence (*NKTR*), arylsulfatse B (*ARSB*), and insulin like growth factor 1 (*IGF1*) (**Fig. 3a**). Some of the markers have expression changes in the same direction in COVID-19 and AD or AD-related pathologies, including *TGFB1*, *GSTM3*, and *NKTR*. Using the PBMC single-cell RNA-sequencing data, we found that prostaglandin-endoperoxide synthase 2 (*PTGS2*) and period circadian regulator 1 (*PER1*) were significantly elevated in monocytes (**Fig. S3**) of severe COVID-19 patients. *PTGS2* expression was also elevated in the bulk PBMC dataset, although not significantly. *PER1* is a circadian clock gene involved in AD [83]. In the CSF, several AD markers were also altered, such as secreted phosphoprotein 1 (*SPP1*), C-X-C motif chemokine ligand 10 (*CXCL10*), and TNF receptor superfamily member 1B (*TNFRSF1B*) (**Fig. 3b**). *TNFRSF1B* showed consistent expression changes in AD or AD-related pathologies, as well as in COVID-19 patient CSF samples. We also found that CXCL10 protein level was increased in CSF of COVID-19 patients [84] (**Fig. 3b**).

To understand the potential pathological consequences of these alterations by SARS-CoV-2 infection, we interrogated the human protein-protein interactome, the ten HCoVs host factor datasets, and the transcriptome data from PBMCs (**Fig. 3c**) of COVID-19 patients and CSF samples of neuro-COVID-19 patients (**Fig. 3d**). We selected three AD blood markers (*TGFB1*, *GSTM3*, and *NKTR*) and three CSF markers (*SPP1*, *CXCL10*, and *TNFRSF1B*) as examples. **Fig. 3c** and **Fig. 3d** show the PPIs among these markers (centered nodes) and their neighbors, which interact with many DEGs or SARS-CoV-2 host factors. For example, NKTR interacts with zinc finger CCH-type containing 18 (ZC3H18) (SARS-CoV-2 host factor), small nuclear interacting protein 1 (SNIP1) (SARS-CoV-1 and SARS-CoV-2 host factor), and casein kinase II subunit alpha (CSNK2A2) (SARS-CoV-1, SARS-CoV-2, and MERS-CoV host factor). NKTR and its PPI partners transcription initiation factor TFIID subunit 1 (TAF1), 40S ribosomal protein S14 (RPS14), and arrestin beta 2 (ARRB2) are differentially expressed in the PBMCs of COVID-19 patients. ARRB2 inhibits toll-like receptor 4 (TLR4)-mediated inflammatory signaling [85], which is activated by the SARS-CoV-2 spike protein [86]. In CSF, innate immune genes *SPP1*, *CXCL10*, and *TNFRSF1B* are differentially expressed in COVID-19 vs. non-COVID-19 patients. Many of their PPI partners are also SARS-CoV-2 host factors, among which some are innate immune gene products, such as integrin subunit beta 1 (ITGB1), which is highly expressed in airway epithelial cells [87], and TNF receptor associated factor 3 (TRAF3), which controls type I interferon (IFN-I) production [88]. Integrins may function as an alternative docking receptor for SARS-CoV-2 [89], and ITGB1 is also essential for migration of monocytes across the endothelium [90].

In summary, expression of these selected AD markers was significantly altered by SARS-CoV-2 infection. Using network and multi-omics data analysis, we found that SARS-CoV-2 infection impacts several immune-related genes/pathways that could lead to AD-like neurologic impairment.

### Elevated expression of SARS-CoV-2 host factors in brain endothelial cells

We next evaluated the susceptibility of brain endothelial cells to SARS-CoV-2 infection and potential microvascular injury. For this, we analyzed the single-nuclei RNA-sequencing dataset from the prefrontal cortex region of 12 AD patients and 9 cognitively healthy controls [42] (**Fig. 4a**). We examined expression of SARS-CoV-2 entry factors across the six cell types: astrocytes, endothelial cells, excitatory neurons, inhibitory neurons, microglia, and oligodendrocytes (**Fig. 4b**). We observed low expression levels of *ACE2*, transmembrane serine protease 2 (*TMPRSS2*), furin (*FURIN*), and neuropilin 1 (*NRP1*) in neurons in both AD patients and healthy controls. For example, *ACE2* and *TMPRSS2* are mostly absent across all six cell types. However, *NRP1* is expressed in endothelial cells, astrocytes, and microglia, and expression is elevated in these cell types than in neurons. NRP1 was reported to mediate SARS-CoV-2 cell entry in addition to ACE2 and TMPRSS2 [91, 92]. Basigin (*BSG*) is much more strongly expressed in endothelial cells than other cell types, and has been reported as a docking receptor for SARS-CoV-2 [93], in addition to ACE2 and NRP1. Among the proteases, *FURIN* has an elevated expression in endothelial cells compared to other cell types, and cystatin B (*CSTB*) is highly expressed in microglia. Differential gene expression analysis confirmed that *BSG* and *FURIN* have significantly higher expression in the brain endothelial cells than in other cell types (**Table S6**). In addition to these SARS-CoV-2 entry factors, we also found elevated expression of antiviral defense system genes in brain endothelial cells, including lymphocyte antigen 6 family member E (*LY6E*), interferon induced transmembrane protein 2 (*IFITM2*) and 3 (*IFITM3*), and interferon alpha and beta receptor subunit 1 (*IFNAR1*). These findings are further confirmed in a second single-nuclei RNA-sequencing dataset [43] (**Fig. S4**). LY6E impairs entry of coronavirus by inhibiting spike protein-mediated membrane fusion [94]. IFN-I receptors (IFNAR) play important roles in IFN-I-mediated antiviral immunity [95], and IFN-induced transmembrane protein 3 (IFITM3) inhibits SARS-CoV-2 cell entry [96, 97]. IFITM3 is also associated with AD through its ability to bind and upregulate γ-secretase, which leads to increased Aβ production [98]. Network analysis also revealed several important PPI partners of these antiviral defense genes (**Fig. 4c**), such as signal transducer and activator of transcription 3 (*STAT3*) and janus kinase 1 (*JAK1*). These immune genes are the HCoVs host factors, and have significantly elevated expression in endothelial cells compared to other cell types of the brain. The JAK-STAT signaling pathway mediates the biological functions of several cytokines involved in cytokine release syndrome (CRS) [99], which is common in COVID-19 [100]. Notably, JAK inhibition reduces SARS-CoV-2 infection in liver and reduces overall morbidity and mortality in COVID-19 patients in a pilot clinical trial [101]. Inhibition of JAK-STAT signaling has therefore been proposed as a treatment strategy for COVID-19 [102].

**Fig. 4.**
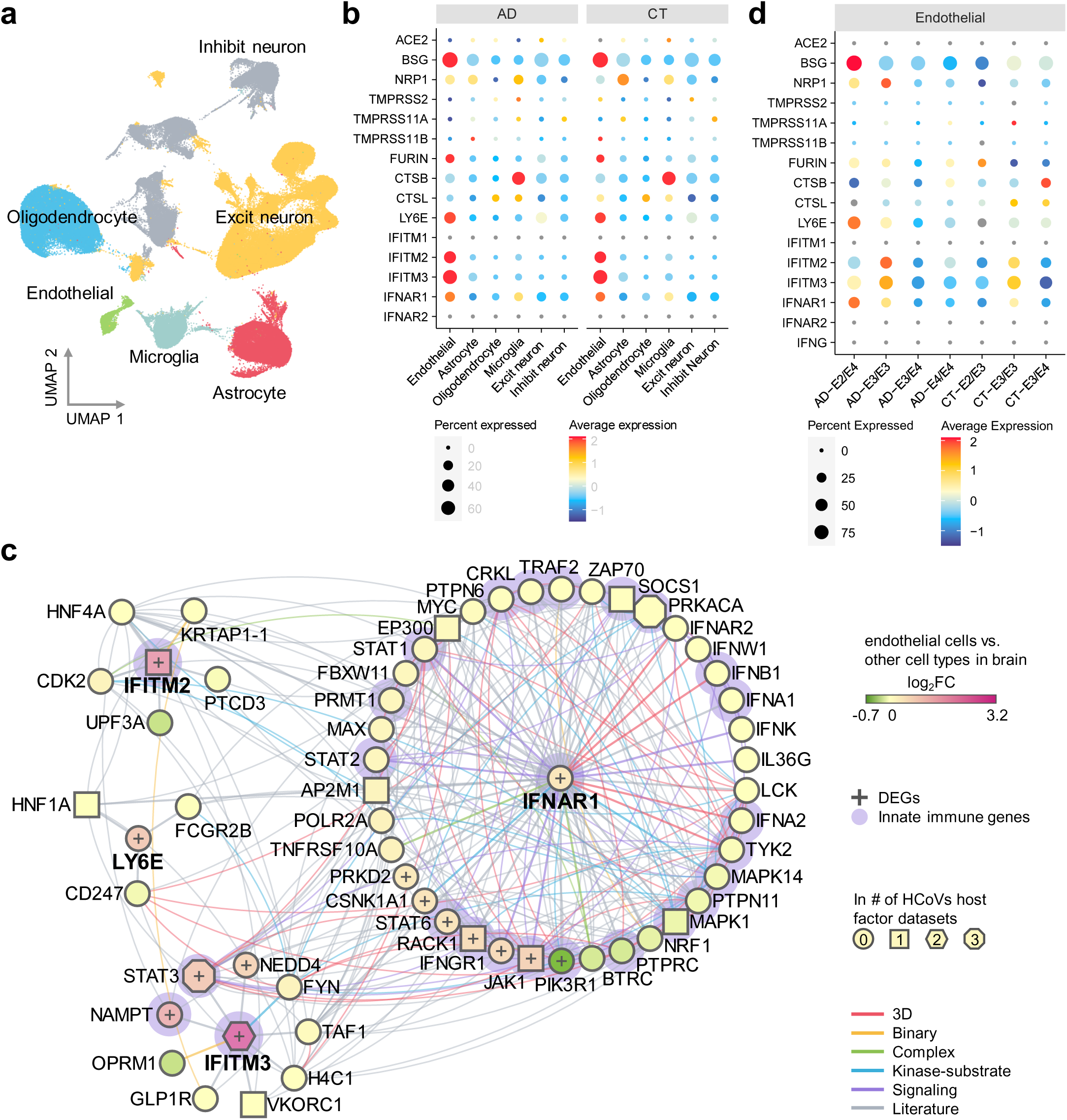
Elevated expression of SARS-CoV-2 host factors in human brain endothelial cells. (**a**) UMAP visualization of the single-nuclei RNA-sequencing dataset from the prefrontal cortex region of Alzheimer’s disease (AD, n=12) patients and healthy controls (CT, n=9). (**b**) Expression of the entry factors and antiviral defense proteins in different cell types in AD and CT groups. (**c**) Network analyses of the antiviral defense genes that are differentially expressed in brain endothelial cells vs. other cell types. Node shape indicates the number of SARS-CoV-2 host factor datasets that contain the node. Edge colors indicate the protein-protein interaction source type. (**d**) Expression of the entry factors and antiviral defense proteins in individuals with different *APOE* genotypes (AD-E3/E3 n=4, AD-E4/E4 n=2, AD-E3/E4 n=5, AD-E2/E4 n=1, CT-E2/E3 n=2, CT-E3/E3 n=5, CT-E3/E4 n=2).

### Lack of expression of antiviral defense genes in *APOE* E4/E4 individuals

It has been suggested that SARS-CoV-2 neurotropism in neurons and astrocytes may be affected by the *APOE* genotype [103]. Individuals carrying *APOE* E2 have decreased AD risk [104, 105], and those carrying *APOE* E4 have increased risk [105], relative to carriers of the normal *APOE* E2 allele. Therefore, we examined expression of these genes in endothelial cells (**Fig. 4d**) and other cell types (**Fig. S5**). Expression of *BSG*, *NRP1*, *FURIN*, and *CTSB* varies by *APOE* genotype. For example, *NRP1* is more highly expressed in E3/E3 AD patients than in E4/E4 AD patients (**Table S7**). Importantly, *LY6E*, *IFITM2*, *IFITM3*, and *IFNAR1* have higher expression in E3/E3 AD patients than in E4/E4 AD patients. These results suggest that AD patients with *APOE* E4/E4 genotype may have a less active antiviral defense system, which could render them at increased risk for SARS-CoV-2 infection.

### Overall low expression of SARS-CoV-2 host factors in human brain

As SARS-CoV-2 infection depends on key entry factors, including *ACE2*, *TMPRSS2*, *FURIN*, and *NRP1*, we first examined expression of these entry factors in healthy tissues using GTEx data [68]. We found overall low expression of SARS-CoV-2 entry factors (*ACE2*, *TMPRSS2*, *FURIN*, and *NRP1*) in the human brain (**Fig. S6**). Brain-specific expression of the four SARS-CoV-2 entry factors (blue bars in the highlighted yellow column of **Fig. 5a**) are lower than in other tissues.

**Fig. 5.**
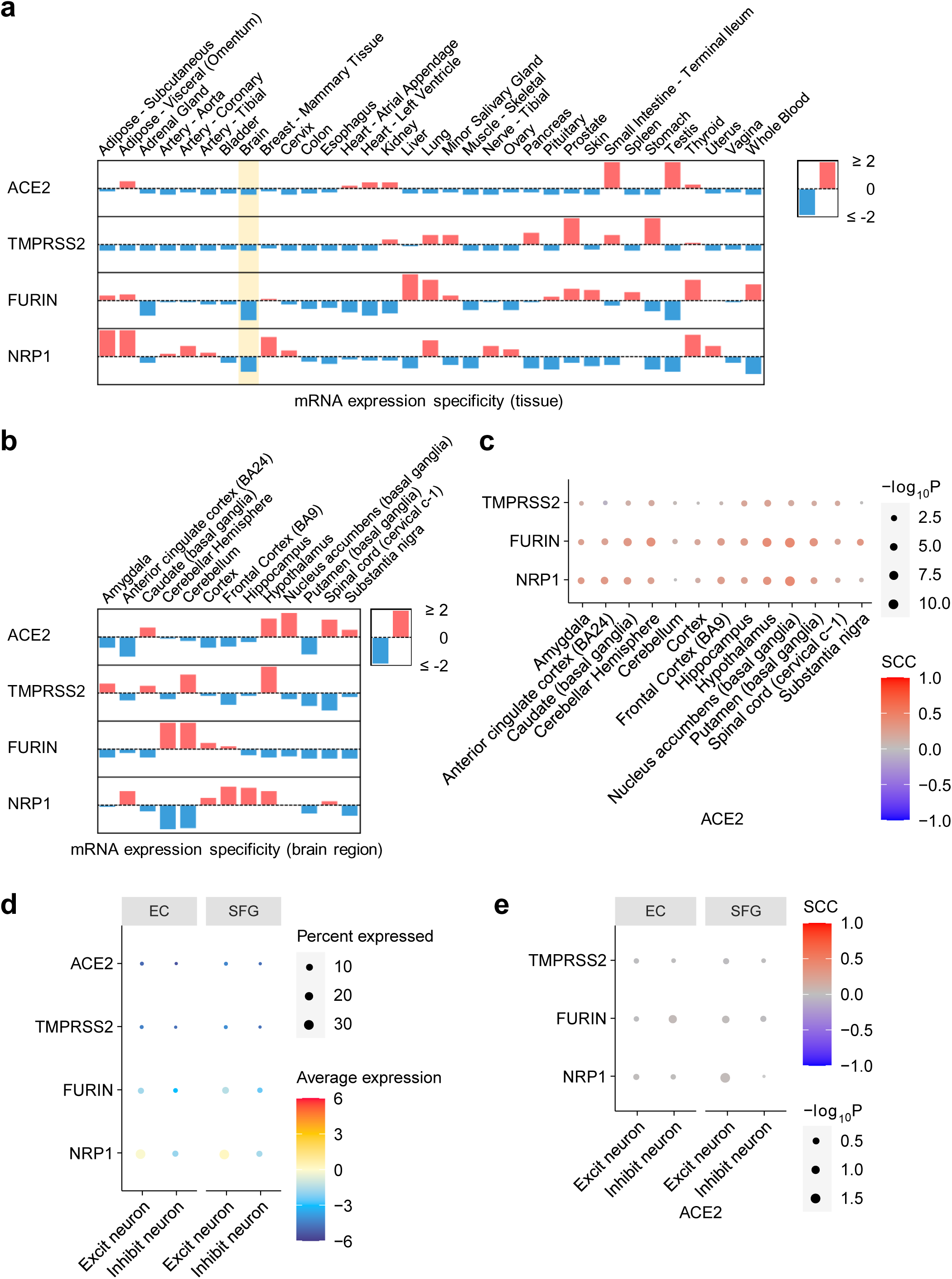
Expression of key SARS-CoV-2 entry factors across 33 human tissues, 13 brain regions, and brain cell types/subpopulations. (**a**) Expression specificity of key SARS-CoV-2 entry factors in 33 tissues and (**b**) expression specificity of these genes in 13 brain regions using data from the GTEx database (see Methods). (**c**) Co-expression of *TMPRSS2, FURIN*, and *NRP1* vs. *ACE2* in the brain regions. (**d**) Expression of key SARS-CoV-2 entry factors in the neuron cells. (**e**) Co-expression of TMPRSS2, FURIN, and NRP1 vs. ACE2 in the neuron. SCC, Spearman’s rank correlation coefficient. EC, entorhinal cortex. SFG, superior frontal gyrus.

It is possible that these entry factors express in certain brain regions, such as thalamus, brain stem, and hippocampus, which may be targeted by SARS-CoV-2 from the olfactory bulb [106, 107]. Therefore, we further examined expression of these entry factors across different brain regions. Among the 13 brain regions, no region showed high specificity for *ACE2*, *TMPRSS2*, *FURIN*, or *NRP1* (**Fig. 5b** and **Fig. S7**). The Spearman’s rank correlation coefficient (ρ) for *TMPRSS2*, *FURIN*, and *NRP1* with *ACE2* does not show a co-expression (|ρ|_max_=0.42 for *ACE2* and *FURIN* in nucleus accumbens) in any of the 13 brain regions (**Fig. 5C**).

It has been reported that ACE2 has an overall low expression in lung [108, 109], as also shown in **Fig. 5a**, but higher expression in certain cell types such as lung alveolar type II (AT2) epithelial cells [108], bronchial secretory cells [110], nasal mucosa [109], and absorptive enterocytes in the ileum [111]. This prompted us to investigate the brain expression of the entry factors at the single-cell/nuclei level. Using single-nuclei RNA-sequencing data of the caudal entorhinal cortex and the superior frontal gyrus from AD patients [41], we examined expression of the four key SARS-CoV-2 entry factors in the excitatory neuron and inhibitory neuron cells (**Fig. 5d**). Notably, we found very low expression of SARS-CoV-2 entry factors as well, consistent with our findings shown in **Fig. 4b**. In addition, co-expression of *TMPRSS2*, *FURIN*, or *NRP1* with *ACE2* is low (**Fig. 5e**, |ρ|_max_=0.03 for *ACE2* and *FURIN* in inhibitory neurons in the entorhinal cortex region). These results suggest that neurons are unlikely to be a direct target for SARS-CoV-2 infection. However, we should note that even though its expression is low overall, *NRP1* has a relatively higher expression than the other three genes. Together, these expression results at the tissue, brain region, and single-nuclei levels suggest that SARS-CoV-2 is unlikely to directly invade brain, and that cognitive impairment with COVID-19 is more likely caused by neuroinflammation (**Fig. 3**) and microvascular injury (**Fig. 4**).

## Discussion

The negative effects of COVID-19 on the CNS may have a long-term impact that could possibly increase the likelihood of developing AD-like dementia [1, 2, 4, 5, 112]. Here, we investigated the potential mechanisms for this effect. Using network proximity measure in the human PPI, we found strong network-based relationship between SARS-CoV-2 host factors (based on PPI assays and CRISPR-Cas9 genetic assays) and disease-associated genes/proteins of dementia-like cognitive impairment. Network analysis of the SARS-CoV-2 host factors and AD-associated genes/proteins reveals that these two sets have significant network proximities in the human interactome. Several AD-associated proteins were highlighted, including RAB7A, TGFB1, and VCAM1, with potentially high impact on the network according to their degrees and eigenvector centralities. In addition, expression of these genes is also altered in COVID-19 patients based on the results of transcriptomic analyses.

Previous studies have shown that SARS-CoV-2 is absent from the brain [12] and CSF [13]. However, evidence also exists that SARS-CoV-2 may directly infect the brain [9–11]. To test the possibility of direct brain invasion by SARS-CoV-2, we investigated the expression of key entry factors of SARS-CoV-2 at three levels: tissue, brain regions, and brain cell types. We found very low expression of *ACE2* and *TMPRSS2* in the brain and neurons. ACE2 is the main known SARS-CoV-2 docking receptor [108–110]; yet, it has little to no expression in neurons (**Fig. 4b** and **Fig. 5d**). Recent studies found two additional SARS-CoV-2 docking receptors, NRP1 [91, 92] and BSG [93]. *BSG*, *NRP1*, and *FURIN* have elevated expression in the endothelial cells in the prefrontal cortex region of both AD patients and healthy controls compared to other brain cell types (**Fig. 4b**). Our results suggest that it is unlikely for SARS-CoV-2 to target neurons directly via ACE2. However, we cannot rule out the possibility that SARS-CoV-2 may enter the brain through the cerebral endothelium using receptors such as BSG and NRP1 or other unknown entry factors. In addition, other HCoVs, including HCoV-229E and HCoV-OC43, have been detected in human brains [113].

Neuroinflammation is a major hallmark of AD, and we analyzed the expression of AD blood and CSF markers in PBMCs and CSF of COVID-19 patients. We identified several AD marker genes (e.g., *NKTR*, *GSTM3*, *TGFB1*, *TNFRSF1B*, *SPP1*, and *CXCL10*) which may provide insights into the shared pathobiology of cognitive dysfunction in COVID-19 and AD. These genes were significantly altered in PBMCs or CSF of COVID-19 patients. Network analysis showed that these genes are enriched in PPIs of immune-related gene products, such as ITGB1 and ARRB2. Moreover, many of the PPI partners of these genes are either the host factors of SARS-CoV-2, or are significantly altered in COVID-19 patients, or both. In addition, the endothelial cells also have elevated expression of antiviral defense genes (*LY6E*, *IFITM2*, *IFITM3*, and *IFNAR1*) (**Fig. 4b**). We identified important PPI partners (*STAT3* and *JAK1*) of these genes using network analysis combined with SARS-CoV-2 host factor datasets and differential expression analyses. Due to the inflammation role of the JAK-STAT signaling pathway in COVID-19, its inhibition by baricitinib has been studied as a potential treatment [102] in several clinical trials (NCT04320277 and NCT04321993). We also found that individuals with *APOE* E4/E4 have lower expression of antiviral defense genes compared to individuals with *APOE* E3/E3, suggesting lack of expression of these genes and potentially an elevated risk of SARS-CoV-2 infection. Human-induced pluripotent stem cell models showed an elevated susceptibility to SARS-CoV-2 infection in *APOE* E4/E4 brain cells [103]. Further observations of *APOE*-related susceptibility to SARS-CoV-2 infection are warranted.

In summary, our observations provide mechanistic insights into two questions: (a) whether SARS-CoV-2 infection could potentially increase the risk of AD and AD-like dementia; and (b) whether individuals with AD and AD-like dementia have increased risk of SARS-CoV-2 infection. Our analyses show a low possibility of direct brain invasion by SARS-CoV-2 (**Fig. 5**). However, we found significant mechanistic overlap between AD and COVID-19 (**Fig. 2**) centered on neuroinflammation and microvascular injury pathways or processes (**Fig. 3** and **Fig. 4**). It was found that dementia patients had twice the risk of COVID-19 compared to those without dementia [6]. Although nursing home stays were adjusted in this study [6], it could still potentially explain the high risk in dementia patients, due to a higher nursing home stay tendency in these patients. We found that the SARS-CoV-2 entry factors and the antiviral defense genes have similar transcriptomic expression in the brain cells between AD patients and control individuals (**Fig. 4b** and **Fig. S4**). These observations do not suggest an elevated risk of COVID-19 in AD patients. Therefore, longitudinal clinical and functional studies are warranted to inspect the causal relationship of dementia and an elevated risk of SARS-CoV-2 infection in the near future.

### Limitations

We acknowledge several limitations. First, our human protein-protein interactome was built using high-quality data from multiple sources; yet it is still incomplete. The PPIs in our interactome is undirected. However, it has been shown that incorporating directionality of the human PPI does not change network proximity results [114]. Therefore, the network associations could be either positive or negative, and require further investigation. In addition, as our network proximity analysis relies on disease-associated genes, literature bias could affect the results because more highly-studied genes are more likely to appear in the dataset. Second, we analyzed expression levels of the key SARS-CoV-2 entry factors and found low expression levels for *ACE2* and *TMPRSS2*. However, we cannot rule out the possibility of SARS-CoV-2 directly targeting the brain via as-yet unidentified mechanisms. Third, possible pathways of neuroinflammation and microvascular injury were tested using data of either individuals with AD or COVID-19, but not both. Future studies using genetics and multi-omics data from individuals with both AD and COVID-19 will be needed to confirm and extend these network-based findings. The significance of our findings in the context of the general population of COVID-19 frequently suffering from “brain fog” without a formal diagnosis of AD needs further investigation.

## Conclusions

In this study, we investigated COVID-19-assoicated neurological manifestations using both network medicine methodologies and bulk/single-cell/single-nuclei transcriptomic data analyses. We identified strong shared neuroinflammatory responses between COVID-19 and AD. Several AD markers (*CXCL10*, *TNFRSF1B*, *SPP1, TGFB1*, *GSTM3*, and *NKTR*) have significantly altered expression in COVID-19 patients. Low expression levels of SARS-CoV-2 entry factors were found in human brains, indicating low possibility of direct brain damage by the virus. Transcriptomic analyses showed elevated expression levels of SARS-CoV-2 host factors (*BSG* and *FURIN*) and antiviral defense genes (*LY6E*, *IFITM2*, *IFITM3*, and *IFNAR1*) in brain endothelial cells compared to other cell types, suggesting possible brain microvascular injury by SARS-CoV-2 infection. In addition, individuals with *APOE* E4/E4 may have increased risk of SARS-CoV-2 infection by loss of expression of antiviral defense genes (*LY6E*, *IFITM2*, *IFITM3*, and *IFNAR1*) compared to individuals with *APOE* E3/E3. Altogether, these results can improve our understanding of COVID-19-associated neurological manifestations and provide guidance for future risk management of potential cognitive impairment by SARS-CoV-2 infection. Our findings could lay the foundation for future research that ultimately leads to testable and measurable serum biomarkers that could identify patients at highest risk of neurological complications with COVID-19.

## Acknowledgements

**Funding**: This work was supported by the National Institute of Aging (R01AG066707 and 3R01AG066707-01S1) and the National Heart, Lung, and Blood Institute (R00HL138272) to F.C. This work has also been supported by the National Institute of Neurological Disorders and Stroke (3R01NS097719-04S1) to F.C. and L.J. This work has also been supported in part by the VeloSano Pilot Program (Cleveland Clinic Taussig Cancer Institute) to F.C.

## Author contributions

F.C. conceived the study. Y.Z., J.X., and Y.H. performed data processing and analyses. A.K., R.M., H.Y., Y.L., J.B.L., A.A.P., and L.J. discussed and interpreted all results. Y.Z. and F.C. wrote and all authors critically revised the manuscript and gave final approval.

## Availability of data and materials

The transcriptomic datasets used in this study (GSE147528, GSE157827, GSE138852, GSE157103, GSE149689, and GSE163005) were downloaded from the NCBI GEO database (https://www.ncbi.nlm.nih.gov/geo/). The GTEx v8 dataset was downloaded from https://gtexportal.org/home/. The human protein-protein interactome and the network proximity code can be found in https://github.com/ChengF-Lab/COVID-19_Map. The AD datasets can be found in https://alzgps.lerner.ccf.org/.

## SUPPORTING INFORMATION

### Supplementary Figures

**Fig. S1.**
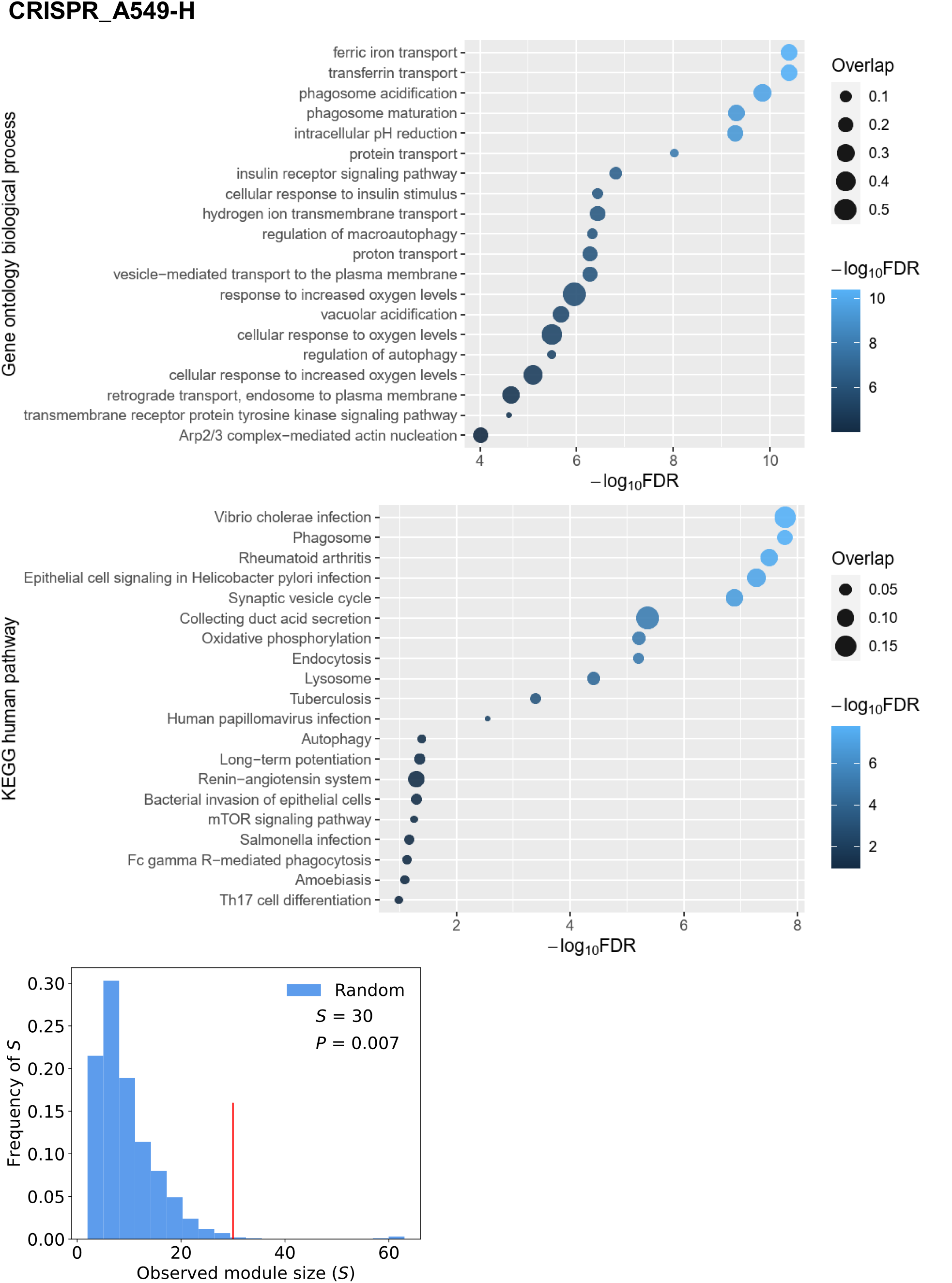
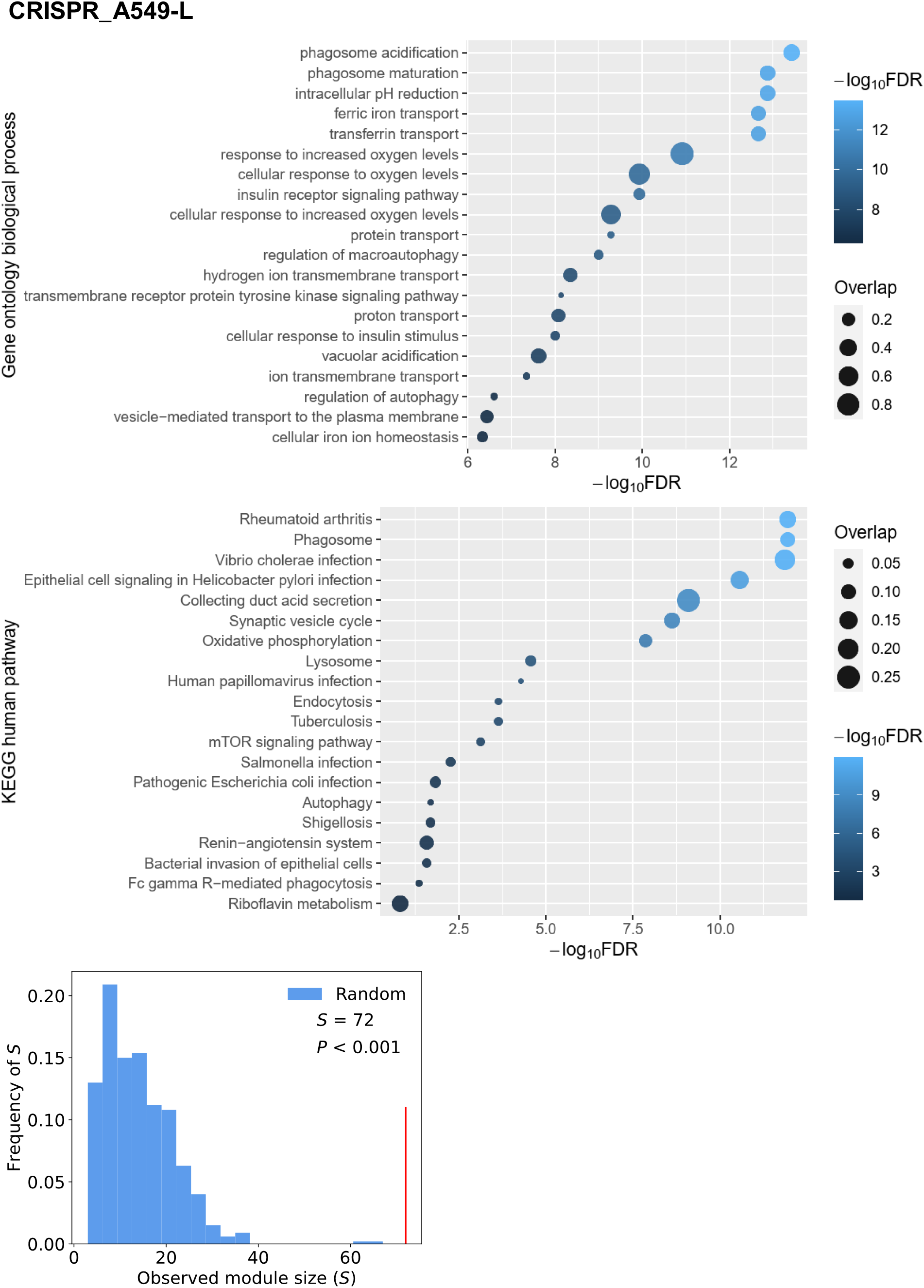
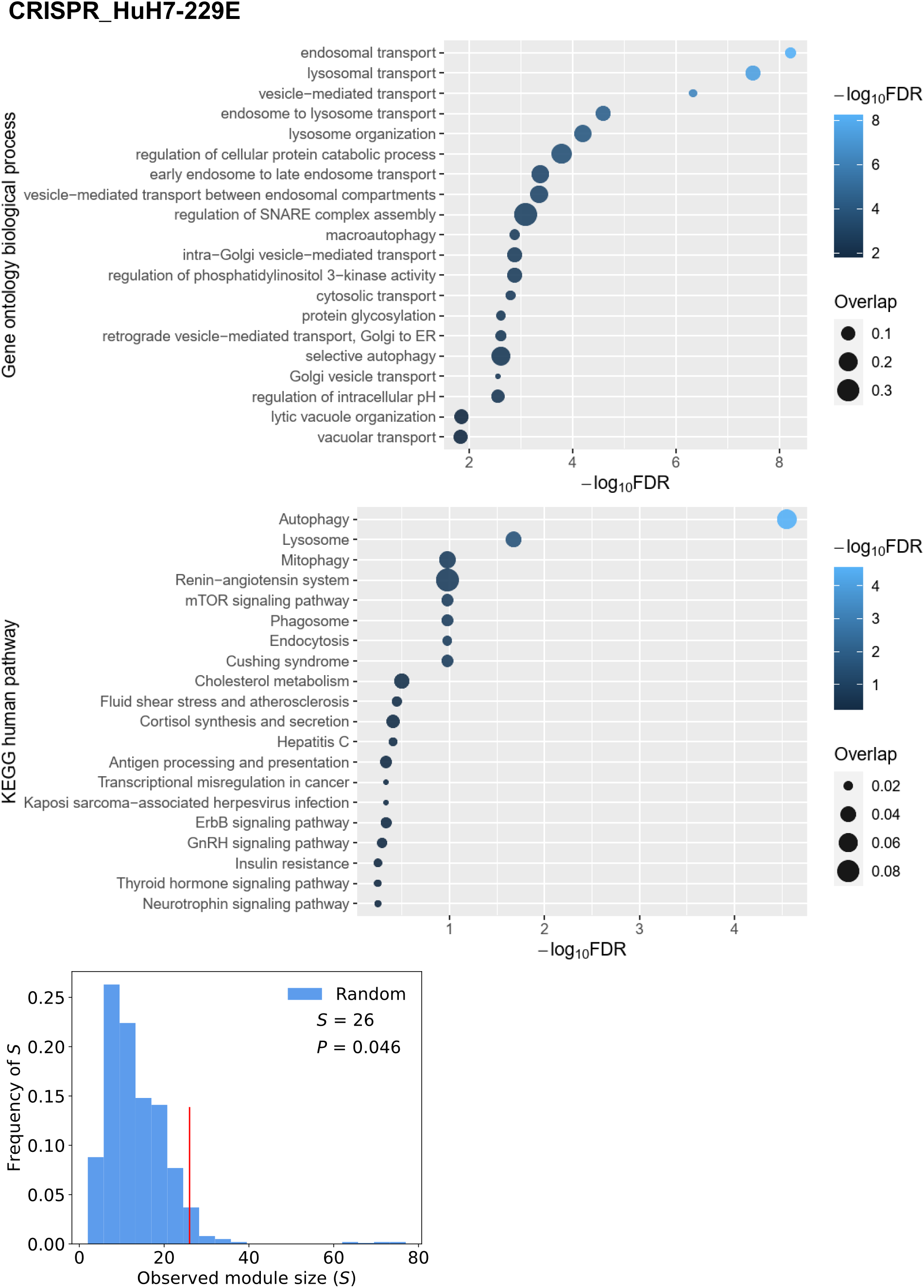
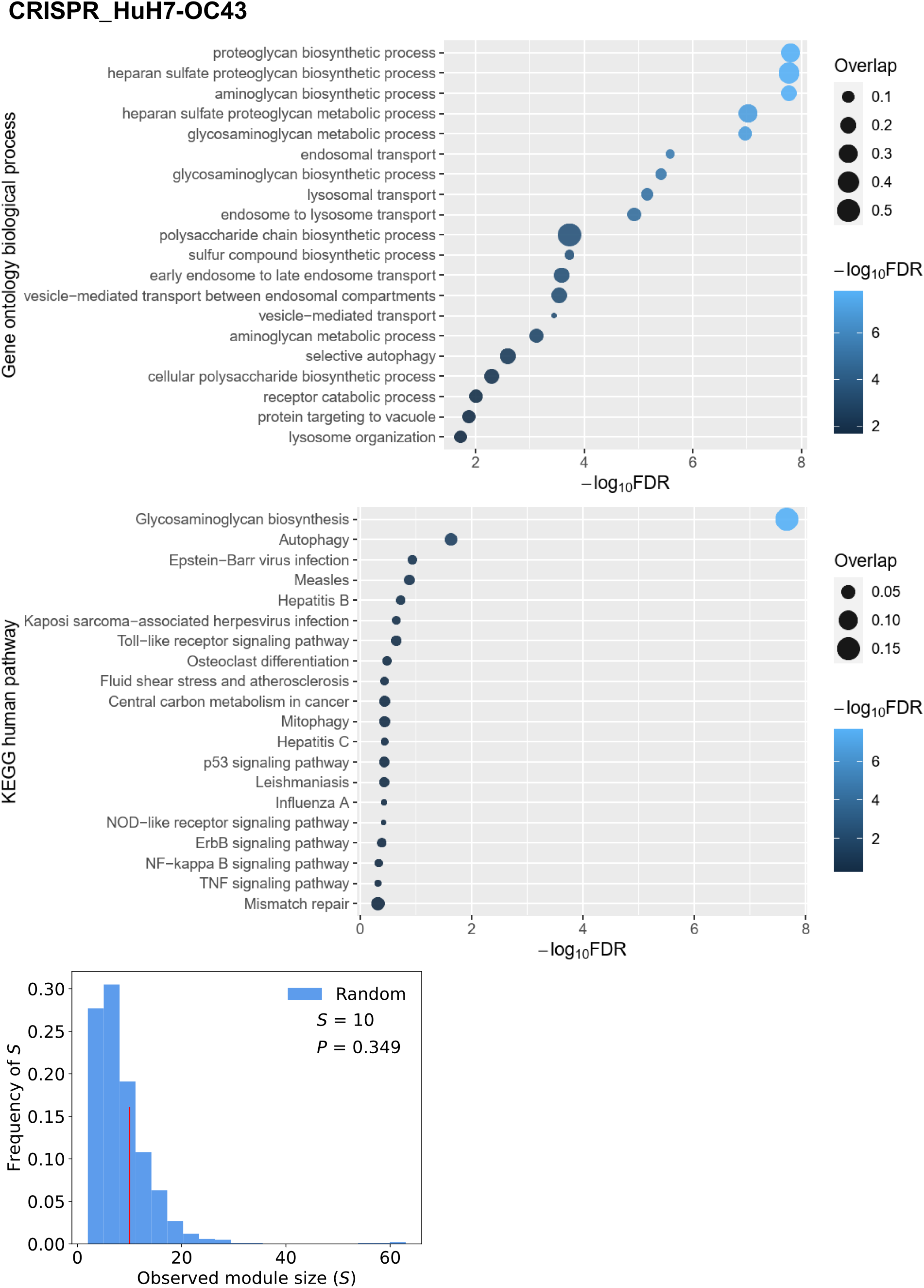
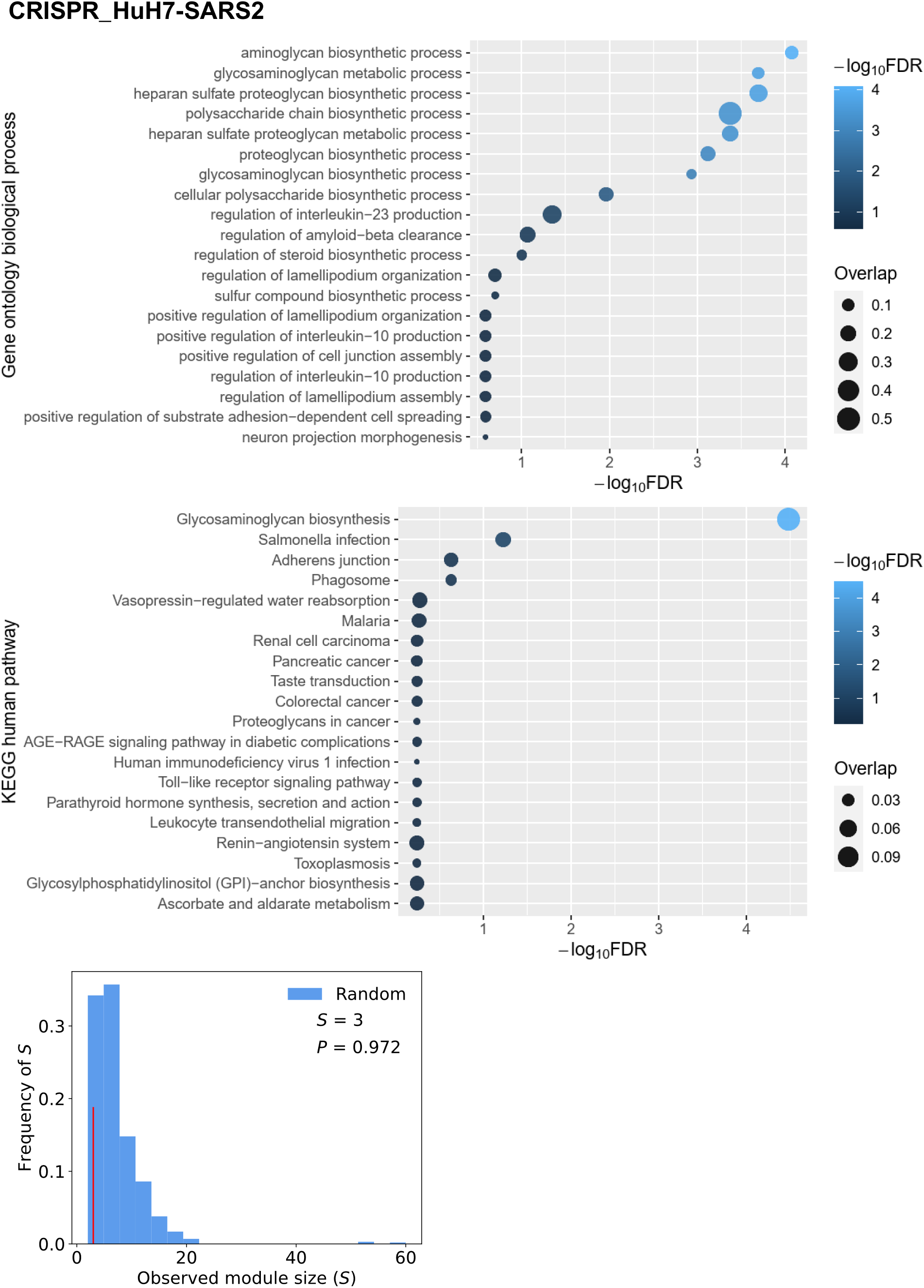
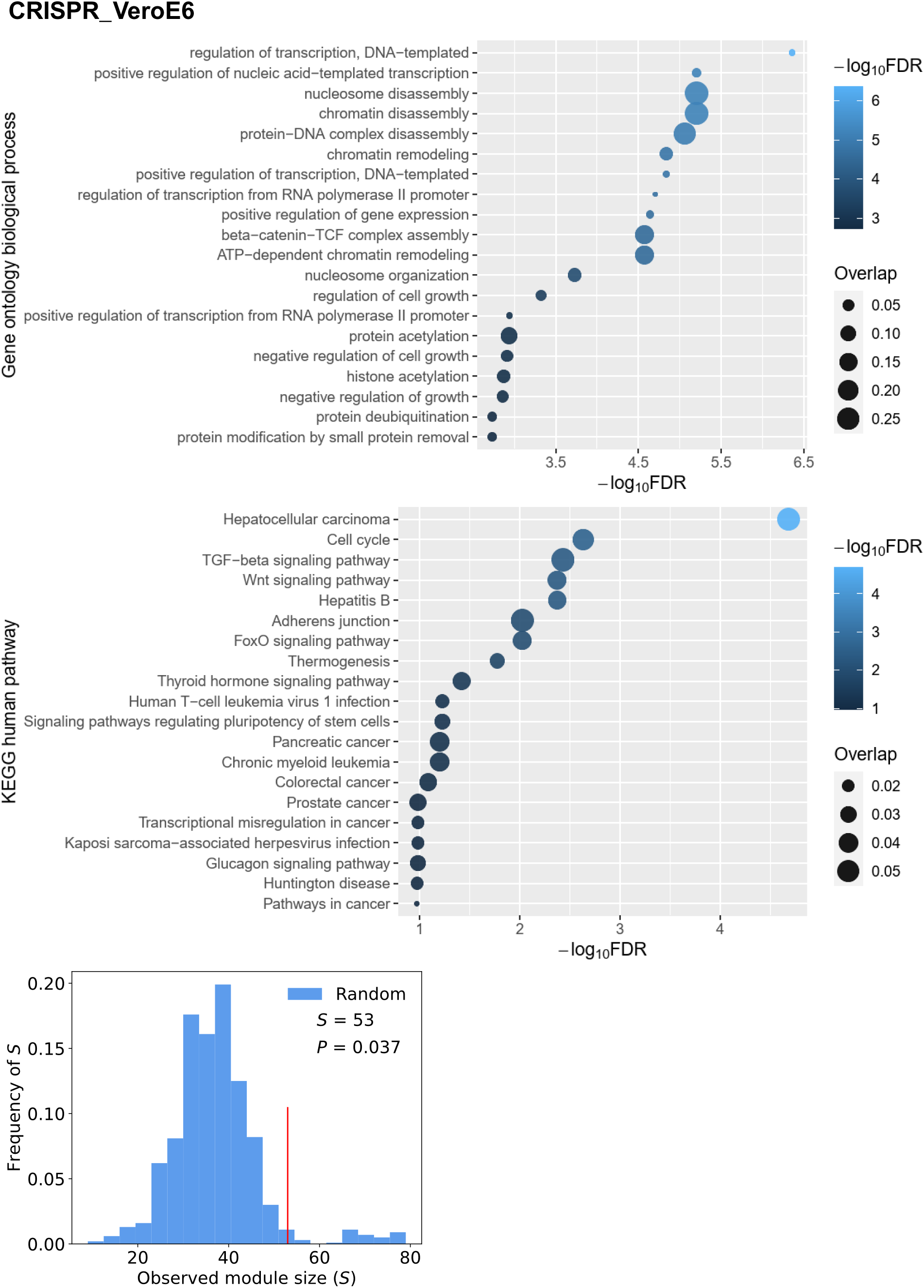
Functional enrichment analysis and largest connected component of the six CRISPR-Cas9-based SARS-CoV-2 host factor datasets. Top 100 genes from each dataset were used for the analyses.

**Fig. S2.**
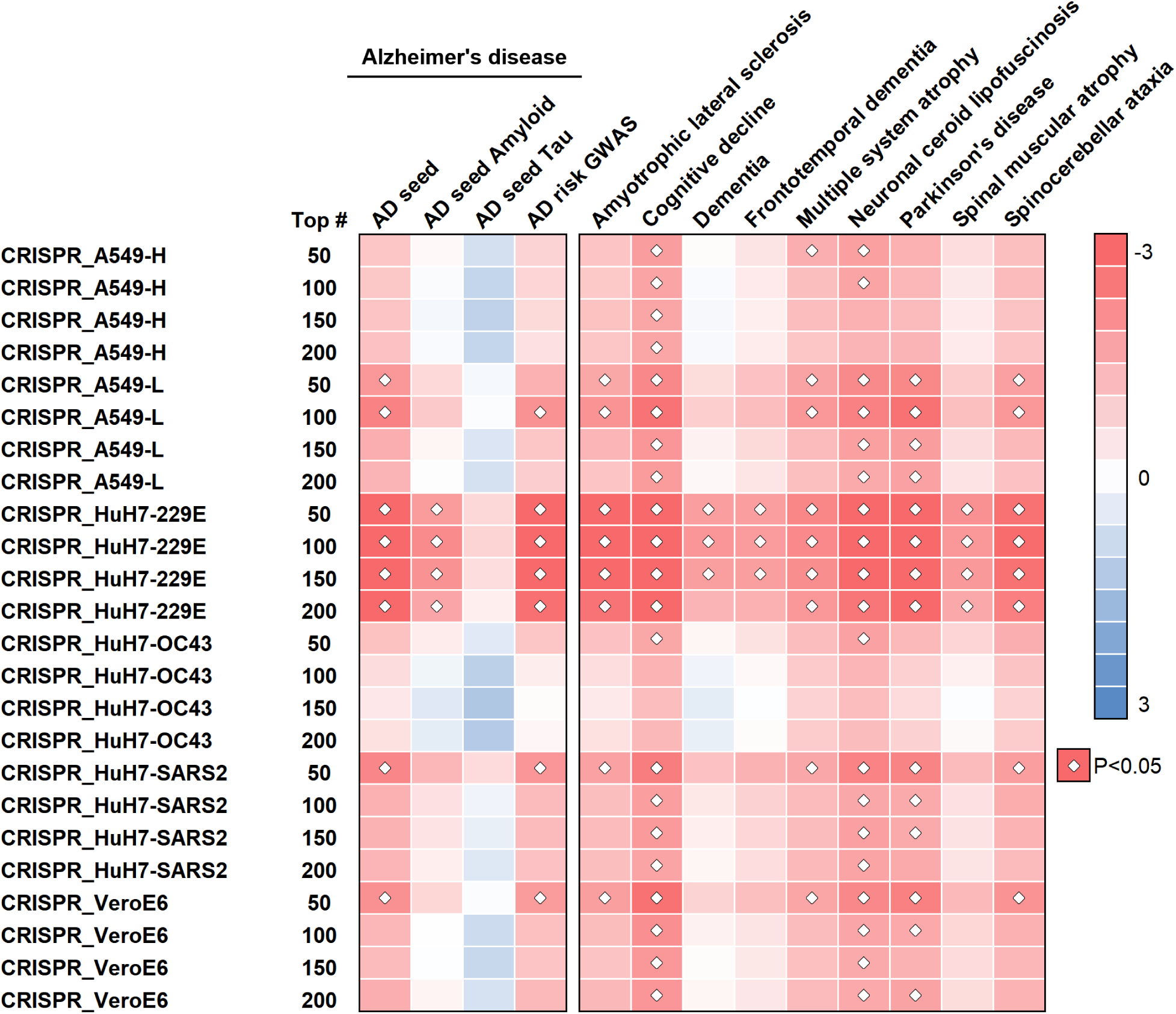
Network proximity results using different numbers of top genes from the CRISPR-Cas9-based SARS-CoV-2 host factor datasets. Heatmap shows the proximities of the CRISPR-Cas9-based SARS-CoV-2 host factor datasets and 10 neurological diseases using different numbers of top genes (i.e., top-50, −100, −150, and −200) from the CRISPR-Cas9 assay.

**Fig. S3.**
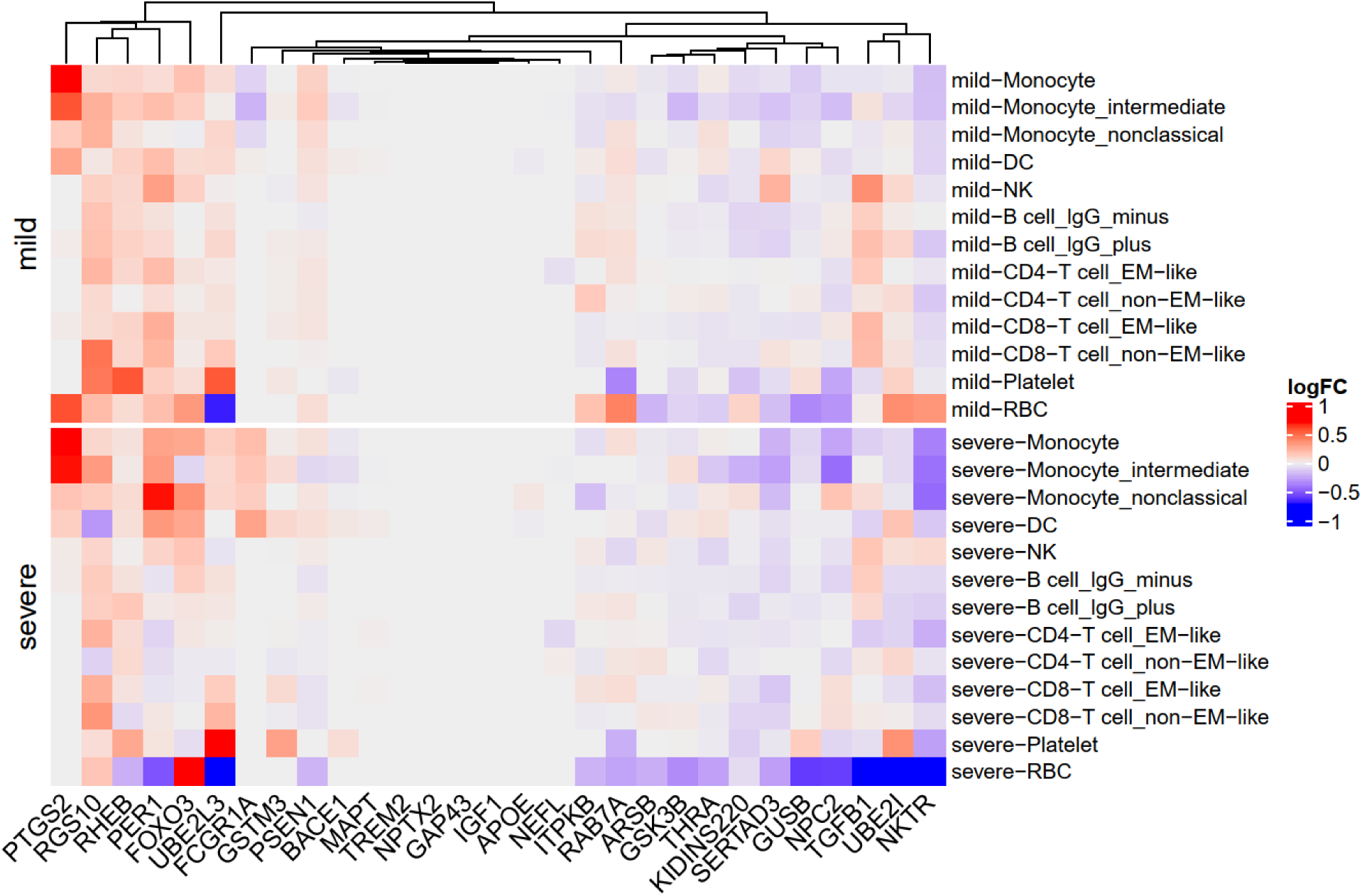
Single-cell level expression of AD blood markers in the PBMC samples of COVID-19 patients. Heatmap shows the expression change in mild / severe COVID-19 patients versus healthy controls. Data source: GSE149689.

**Fig. S4.**
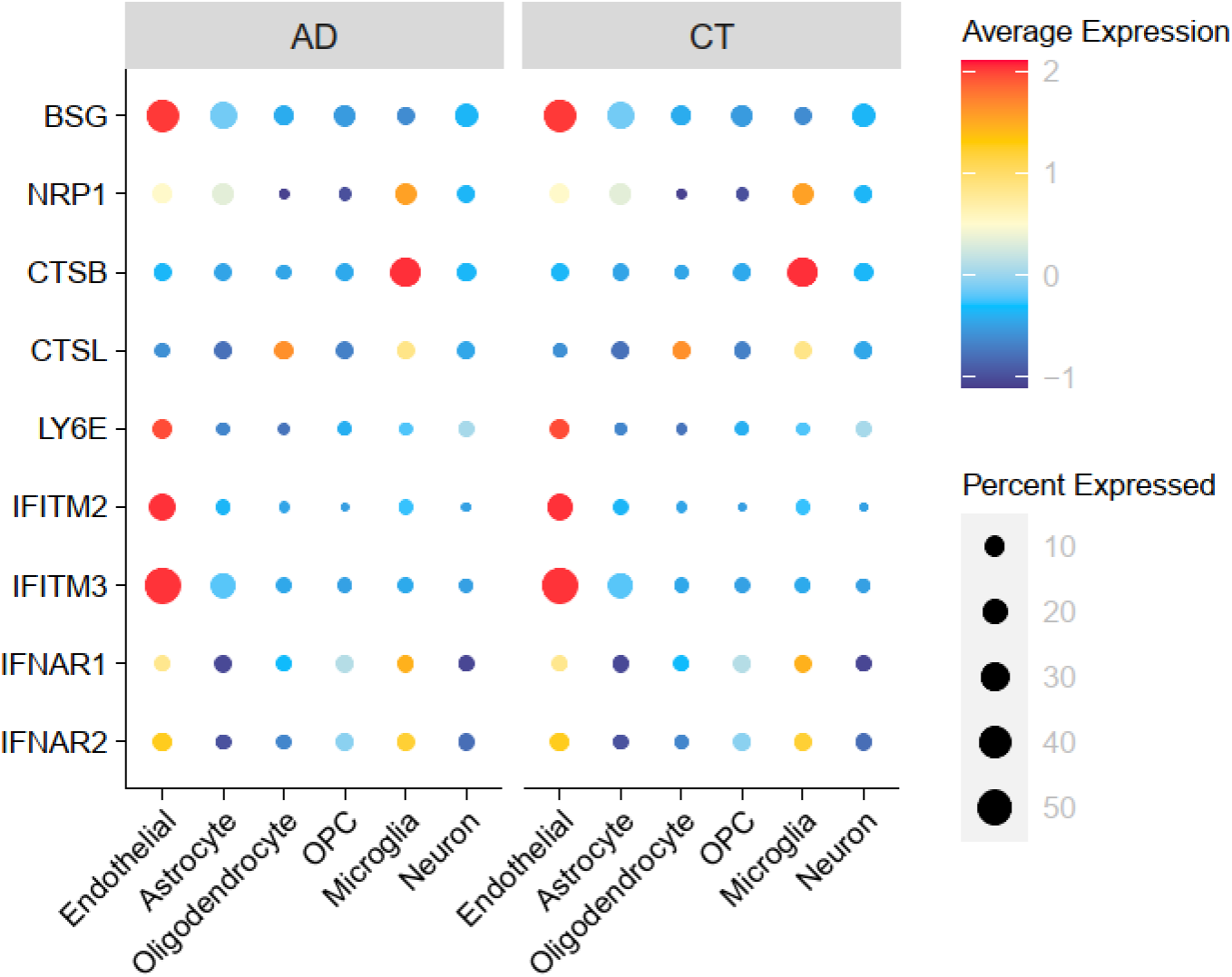
Expression spectrum of the SARS-CoV-2 entry factors in the entorhinal cortex from Alzheimer’s disease patients and controls. AD, Alzheimer’s disease patients. CT, controls. OPC, oligodendrocyte progenitor cell. Data source: GSE138852.

**Fig. S5.**
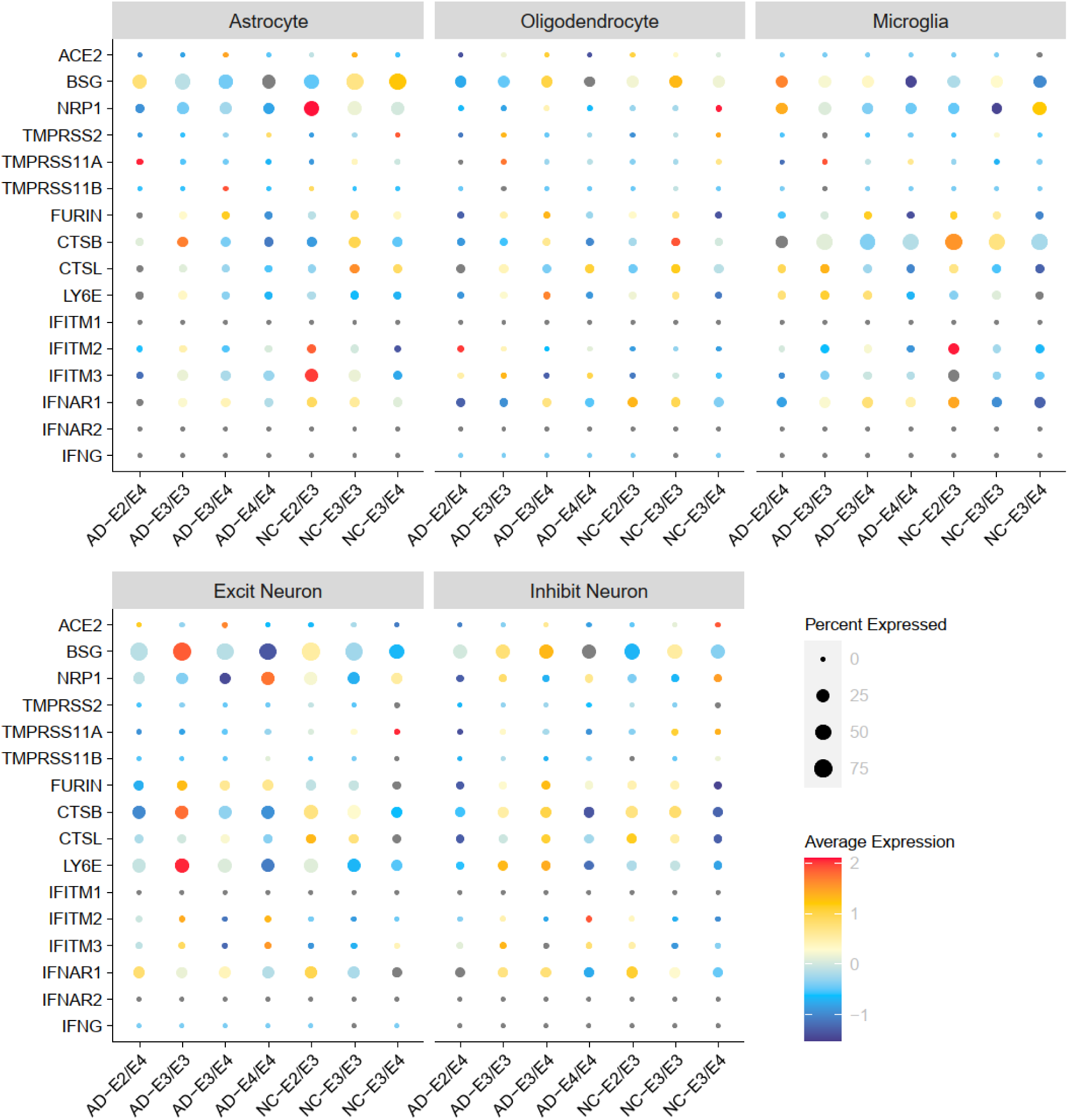
Expression spectrum of the SARS-CoV-2 entry factors in individuals with different *APOE* genotypes. AD, Alzheimer’s disease patients. NC, normal controls. Data source: GSE157827.

**Fig. S6.**
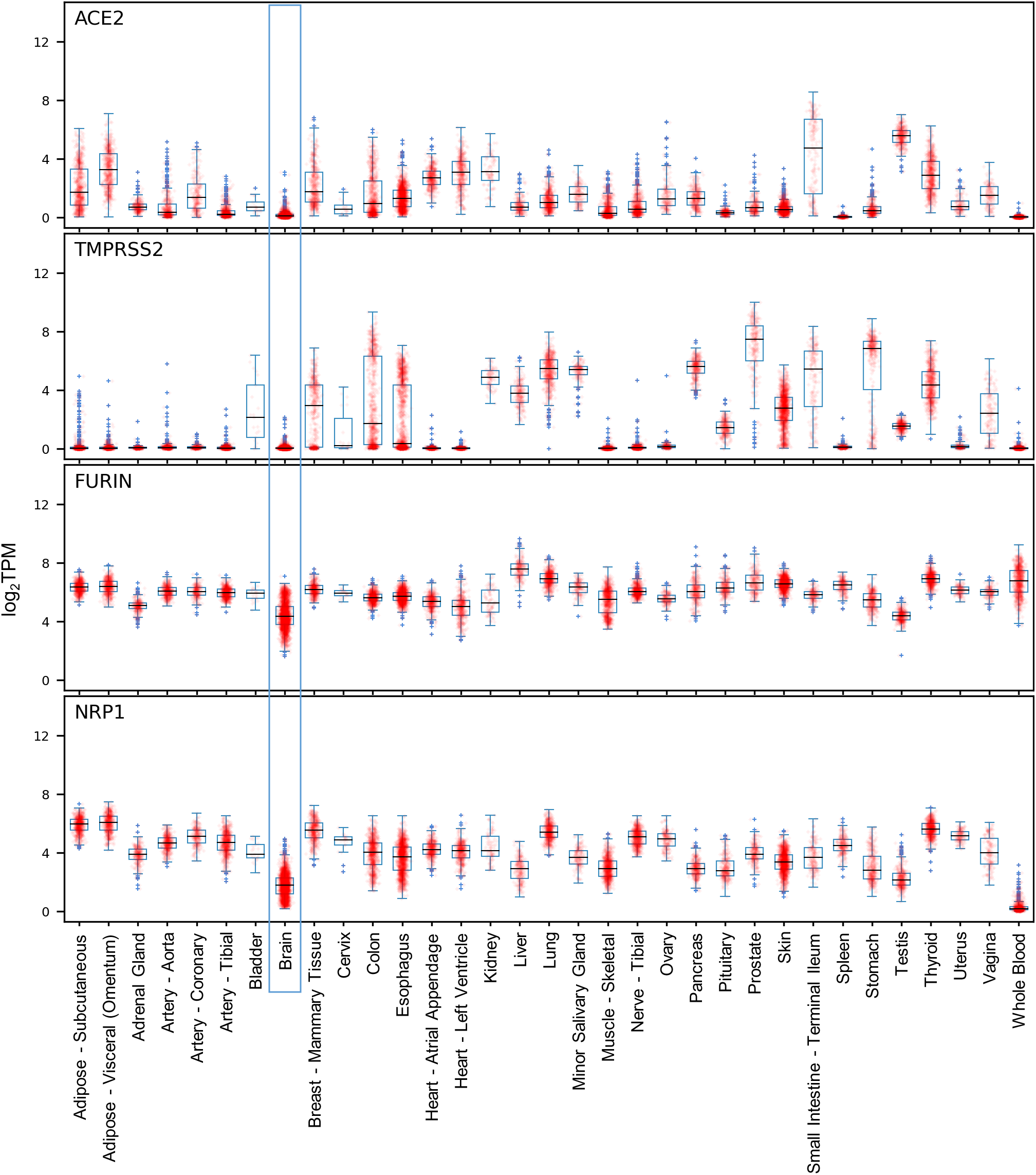
Expression of the key SARS-CoV-2 entry factors in different tissues. Data source: GTEx v8.

**Fig. S7.**
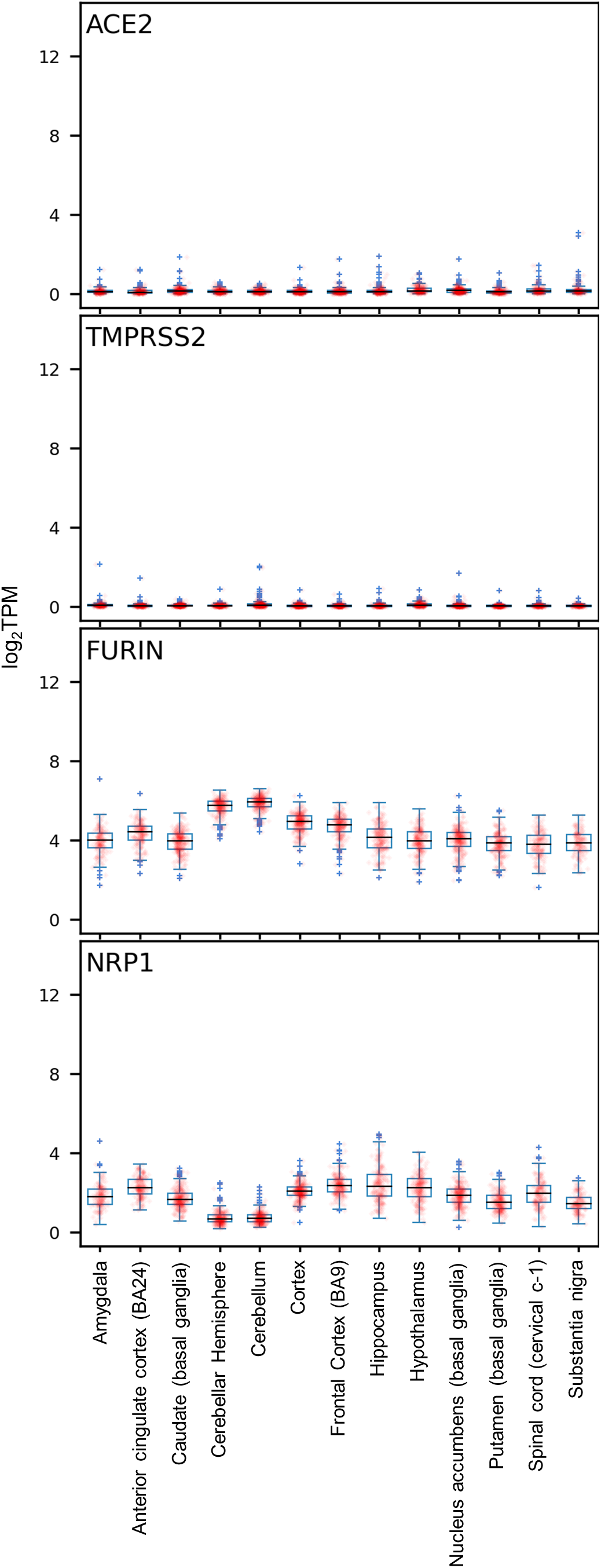
Expression of the key SARSCoV-2 entry factors in different brain regions. Data source: GTEx v8.

### Supplementary Tables

**Table S1.** SARS-CoV-2 host factor datasets. (.xlsx)

**Table S2.** Neurological diseases-associated genes/proteins. (.xlsx)

**Table S3.** Alzheimer’s disease markers and their expressions. (.xlsx)

**Table S4.** Transcriptomic datasets used in this study

| GEO ID | Type | Organism | Sample / Brain region | Groups | Cell types |
| --- | --- | --- | --- | --- | --- |
| GSE147528 | single-nuclei RNA-seq | Homo sapiens | superior frontal gyrus and entorhinal cortex | 10 males with varying stages of Alzheimer's disease (AD) | astrocytes, excitatory neurons, inhibitory neurons, and microglia |
| GSE157827 | single-nuclei RNA-seq | Homo sapiens | prefrontal cortex | 12 AD patients and 9 normal controls | astrocytes, endothelial cells, excitatory neurons, inhibitory neurons, microglia, and oligodendrocytes |
| GSE138852 | single-nuclei RNA-seq | Homo sapiens | entorhinal cortex | AD (n = 6) and healthy controls (n = 6) | astrocytes, endothelial cells, neurons, microglia, oligodendrocytes, and oligodendrocyte progenitor cells |
| GSE157103 | bulk RNA-seq | Homo sapiens | peripheral blood mononuclear cell (PBMC) | 66 intensive care unit (ICU) patients (COVID-19 patients n = 50 vs. non-COVID-19 patients n = 16), 59 non-ICU patients (COVID-19 patients n = 49 vs. non-COVID-19 patients n = 10), and all 125 patients | N/A |
| GSE149689 | single-cell RNA-seq | Homo sapiens | PBMC | 6 samples from severe COVID-19 patients, 4 samples from mild COVID-19 patients, and 4 samples from healthy controls | IgG<sup>-</sup> B cells, IgG<sup>+</sup> B cells, CD4<sup>+</sup> T cell effector memory (EM)-like cells, CD4<sup>+</sup> T cell non-EM-like cells, CD8<sup>+</sup> T cell EM-like cells, CD8<sup>+</sup> T cell non-EM-like cells, dendritic cells, monocytes, intermediate monocytes, nonclassical monocytes, natural killer cells, platelets, and red blood cells |
| GSE163005 | single-cell RNA-seq | Homo sapiens | Cerebrospinal fluid | 8 COVID-19 patients, 9 multiple sclerosis patients, 9 idiopathic intracranial hypertension patients, and 5 viral encephalitis patients | T cells, dendritic cells, and monocytes |

**Table S5.** Raw data and network analysis results of the nodes in Fig. 2b. (.xlsx)

**Table S6.** Differentially expressed genes in brain endothelial cells vs. other cell types. (.xlsx)

**Table S7.** Differentially expressed genes in brain endothelial cells by comparing *APOE* genotype E3/E3 and E4/E4 in Alzheimer’s disease patients. (.xlsx)

## Supplementary Results

We compiled ten SARS-CoV-2 and other HCoVs host factor profiles, including six datasets from CRISPR-Cas9 assays (CRISPR_A549-H, CRISPR_A549-L, CRISPR_HuH7-229E, CRISPR_HuH7-OC43, CRISPR_HuH7-SARS2, and CRISPR_VeroE6), and four datasets for virus-human PPIs (SARS2-PPI, SARS1-PPI, MERS-PPI, and HCoV-PPI) (see Methods). The six CRISPR-Cas9-based datasets adopted genome-scale CRISPR loss-of-function screening methods in the SARS-CoV-2 infected cell lines (as indicated in the dataset name) to identify host factors required for the infection.

As we hypothesized that the SARS-CoV-2 host factors form a subnetwork within the comprehensive human protein interactome, we first computed the largest connected components (LCC) of the CRISPR-Cas9-based datasets. LCC quantifies the number of genes/proteins in the largest subnetwork formed by a dataset. We found that three of these datasets, including CRISPR_A549-H, CRISPR_A549-L, and CRISPR_HuH7-229E, consistently show significantly large LCC (**Table S2**), when we used top-50, −100, and −150 genes. Top-100 revealed the highest number of significant LCCs for the SARS-CoV-2 datasets (CRISPR_A549-H p = 0.007, CRISPR_A549-L p < 0.001, CRISPR_VeroE6 p = 0.037, permutation test, **Table S2**, **Fig. S1**). Therefore, we selected top-100 genes from these datasets for downstream analyses. These results suggest that these datasets form disease modules in the human protein interactome and offer opportunities for network-based discoveries.

Next, we performed functional enrichment analyses for these datasets (**Fig. S1**). We identified several common pathways and GO terms that are enriched in more than three datasets, including autophagy, lysosome, vesicle-mediated transport, endosomal transport, intracellular pH reduction, macromolecule catabolic process, regulation of lysosomal lumen pH, cytosolic transport, and selective autophagy. These datasets also have different functional enrichment. For example, CRISPR_VeroE6 is enriched in functions related to cell cycle, cell growth, and chromatin remodeling, and CRISPR_HuH7-SARS2 is enriched in heparan sulfate biosynthetic functions. These results suggest that the SARS-CoV-2 host factors participate in various essential cellular functions. In addition, these datasets contain complementary information of the cellular states of the SARS-CoV-2 infection and host response.

